# Extracting the Dynamics of Behavior in Decision-Making Experiments

**DOI:** 10.1101/2020.05.21.109678

**Authors:** Nicholas A. Roy, Ji Hyun Bak, The International Brain Laboratory, Athena Akrami, Carlos D. Brody, Jonathan W. Pillow

## Abstract

Understanding how animals update their decision-making behavior over time is an important problem in neuroscience. Decision-making strategies evolve over the course of learning, and continue to vary even in well-trained animals. However, the standard suite of behavioral analysis tools is ill-equipped to capture the dynamics of these strategies. Here, we present a flexible method for characterizing time-varying behavior during decision-making experiments. We show that it successfully captures trial-to-trial changes in an animal’s sensitivity to not only task-relevant stimuli, but also task-irrelevant covariates such as choice, reward, and stimulus history. We use this method to derive insights from training data collected in mice, rats, and human subjects performing auditory discrimination and visual detection tasks. With this approach, we uncover the detailed evolution of an animal’s strategy during learning, including adaptation to time-varying task statistics, suppression of sub-optimal strategies, and shared behavioral dynamics between subjects within an experimental population.

## Introduction

The decision-making behavior of animals in carefully designed tasks is the foundation on which much neuroscience research is built (Carandini, 2012; Krakauer et al., 2017). Many results rely on the accurate and stable decision-making of experimental subjects, whereas other conclusions derive from quantifying specific deficits in choice behavior. For example, causal manipulations of neural activity that result in impaired task accuracy can be traced back to specific disruptions in the sensory processing of task stimuli (Licata et al., 2017; Pisupati et al., 2019). Conversely, manipulations which were found to selectively suppress the impact of task-irrelevant covariates lead to improved accuracy (Akrami et al., 2018). Accurate quantification of decision-making behavior is prerequisite to drawing conclusions from patterns, changes, or abnormalities found in that behavior.

This quantification of the strategies driving an animal’s choices is often challenging for two main reasons. First, an accurate description of behavior needs to not only account for the impact of task-relevant features (e.g., task stimuli), but also measure the influence of other, task-irrelevant covariates (e.g., the animal’s choice on the previous trial). Detecting and disentangling the influence of these variables on an animal’s choices is non-trivial. Is the subject effectively utilizing the task stimuli? In what ways, and to what extent, is trial history affecting decisions?

Second, decision-making behavior is dynamic (Usher et al., 2013; Brunton et al., 2013; Piet et al., 2018). An accurate description of an animal’s behavioral strategy during one session may be inaccurate by the next session. Strategies may evolve significantly within a single session. If decision-making is a product of the influence of many task covariates, as stated above, then the goal of quantifying behavior is complicated by having those influences evolve over time. If task accuracy suddenly drops, has the animal ceased attending to task stimuli, or is the emergence of new, sub-optimal strategy to blame?

These challenges are particularly salient in animal training, especially when animals are first learning a new task (Carandini and Churchland, 2013). Learning necessitates changes in behavior, frustrating conventional analyses that are better suited to the relatively stable behavior seen post-training. Furthermore, behavior during early training is notoriously diverse: different subjects within the same population may utilize entirely different strategies upon encountering a novel task (Cohen and Schneidman, 2013). All research dependent upon animals consistently making accurate decisions must endure the poorly understood and resource-hungry bottleneck of animal training.

Conventional tools for characterizing choice behavior include the psychometric curve and logistic regression, in combination with coarse performance statistics (see, for example, Akrami et al. (2018) and IBL et al. (2020), the sources of the mouse, rat, and human behavior analyzed below) (Green and Swets, 1966). These approaches aggregate data from a range of trials, often spanning many sessions. This collapse across time leaves these approaches ill-equipped to describe behavior that is time-varying, as seen during learning. Although a sizeable literature has focused on the theoretical principles (Sutton, 1988; Courville et al., 2006; Sutton and Barto, 2018) and the biological substrates of learning in the brain (Schultz et al., 1997; Dayan and Balleine, 2002; O’Doherty et al., 2003; Daw and Doya, 2006), there has been a comparative lack of methods for characterizing the dynamics of behavior from experimental datasets.

Several previous studies have nevertheless addressed the topic of dynamic behavior. Smith et al. (2004) introduced a dynamic analysis focused primarily on learning curves, in particular the problem of reliably identifying the time at which learning can be said to occur. While their state-space smoothing algorithm improved over traditional trial-averaged learning curves, the work did not provide a description of detailed changes in behavioral strategy (Suzuki and Brown, 2005; Prerau et al., 2009). Other work from Kattner et al. (2017) extended the standard psychometric curve to allow its parameters to vary continuously across trials. Bak et al. (2016) described a model for defining smoothly evolving weights that could track changing sensitivities to specific behavioral covariates. While the model could track behavioral dynamics in theory, the optimization procedure strongly constrained both the complexity of the model and the size of the data to which it could be applied (Bak and Pillow, 2018).

Here we address these shortcomings by presenting a flexible and efficient method for inferring time-varying behavioral policies in decision-making experiments, facilitated by the statistical innovations presented in Roy et al. (2018a). Animal behavior is quantified at single-trial resolution, allowing for intuitive visualization of learning dynamics and direct analysis of psychophysical weight trajectories. We apply our model to behavioral data collected from throughout the training process of two experimental paradigms, spanning three species (mouse, rat, and human). After validating on simulated data, we leverage our model to analyze an example mouse that learns to synchronize its choice behavior to the block structure of a visual detection task (IBL et al., 2020). We then uncover the dominance of trial history information in rats, but not humans, during early training on an auditory parametric working memory task (Akrami et al., 2018).

An implementation of the model is publicly available as the Python package PsyTrack (Roy et al., 2018b). Furthermore, the Methods include a link to a Google Colab notebook which allows anyone to precisely reproduce all figures directly from the publicly available raw data. We expect that our method will provide immediate practical benefit to animal trainers, in addition to giving unprecedented insight into the development of behavioral strategies.

## Results

Our primary contribution is a method for characterizing the evolution of animal decision-making behavior on a trial-to-trial basis. Our approach consists of a dynamic Bernoulli generalized linear model (GLM), defined by a set of smoothly evolving psychophysical weights. These weights characterize the animal’s decision-making strategy at each trial in terms of a linear combination of available task variables. The larger the magnitude of a particular weight, the more the animal’s decision relies on the corresponding task variable. Learning to perform a new task therefore involves driving the weights on “relevant” variables (e.g., sensory stimuli) to large values, while driving weights on irrelevant variables (e.g., bias, choice history) toward zero. However, classical modeling approaches require that weights remain constant over long blocks of trials, which precludes tracking of trial-to-trial behavioral changes that arise during learning and in non-stationary task environments. Below, we describe our modeling approach in more detail.

### Dynamic Psychophysical Model for Decision-Making Tasks

Although our method is applicable to any binary decision-making task, for concreteness we introduce our method in the context of the task used by the International Brain Lab (IBL) (illustrated in Figure 1A) (IBL et al., 2020). In this visual detection task, a mouse is positioned in front of a screen and a wheel. On each trial, a sinusoidal grating (with contrast values between 0 and 100%) appears on either the left or right side of the screen. The mouse must report the side of the grating by turning the wheel (left or right) in order to receive a water reward (see Methods for more details).

**Figure 1.**
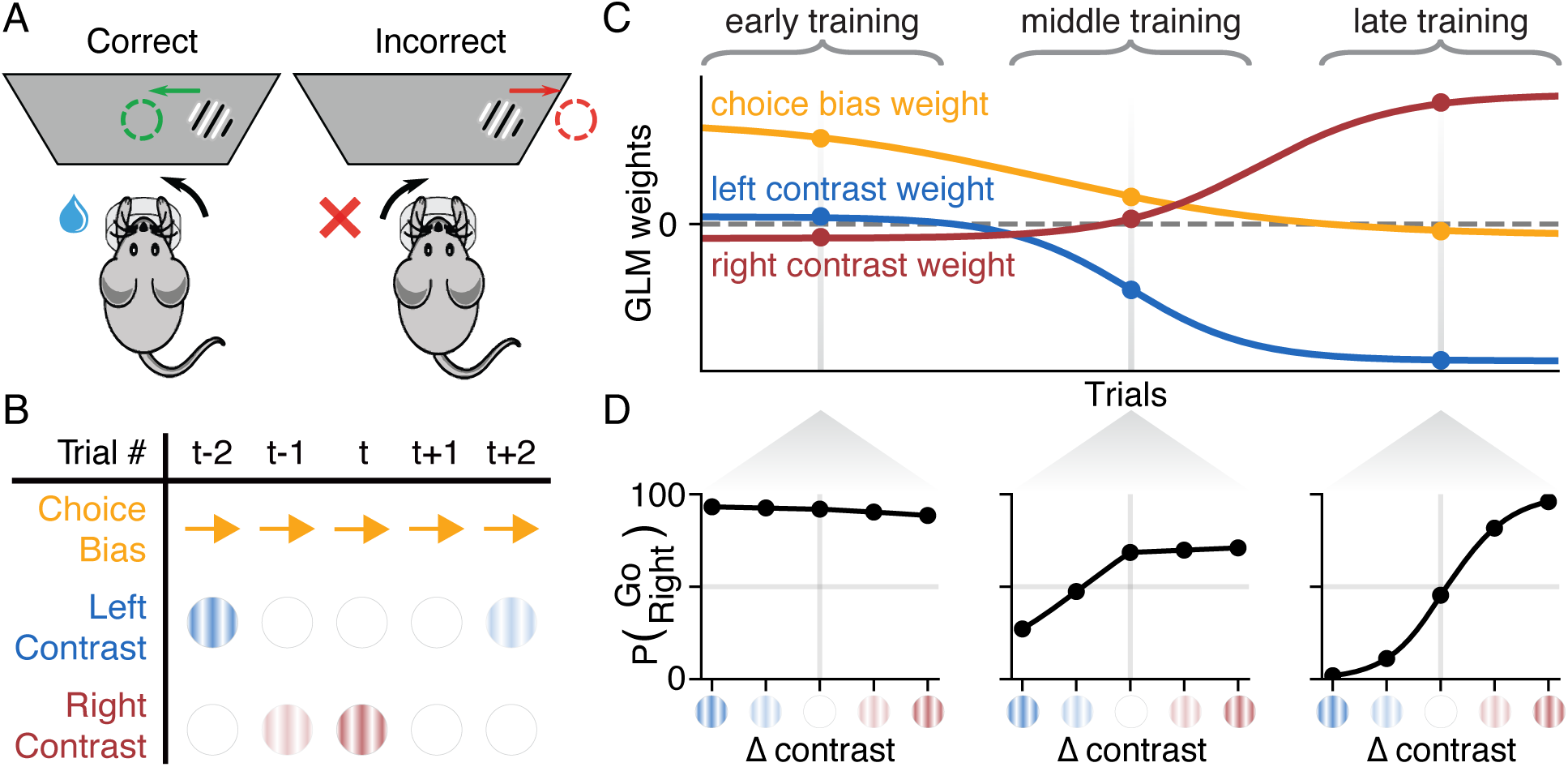
Schematic of Binary Decision-Making Task and Dynamic Psychophysical Model. **(A)** A schematic of the IBL sensory decision-making task. On each trial, a sinusoidal grating (with contrast values between 0 and 100%) appears on either the left or right side of the screen. Mice must report the side of the grating by turning a wheel (left or right) in order to receive a water reward (see Methods for details) (IBL et al., 2020). **(B)** An example table of the task variables **X** assumed to govern behavior for a subset of trials {*t* − 2, *…, t* + 2}, consisting here of a choice bias (a constant rightward bias, encoded as “+1” on each trial), the contrast value of the left grating, and the contrast value of the right grating. **(C)** Hypothetical time-course of a set of psychophysical weights **W**, which evolve smoothly over the course of training. Each weight corresponds to one of the *K* = 3 variables comprising **x**_*t*_, such that a weight’s value at trial *t* indicates how strongly the corresponding variable is affecting choice behavior. **(D)** Psychometric curves defined by the psychophysical weights **w**_*t*_ on particular trials in early, middle, and late training periods, as defined in (C). Initial behavior is highly biased and insensitive to stimuli (“early training”). Over the course of training, behavior evolves toward unbiased, high-accuracy performance consistent with a steep psychometric function (“late training”).

Our modeling approach assumes that on each trial the animal receives an input **x**_*t*_ and makes a binary decision *y*_*t*_ ∈ {0, 1}. Here, **x**_*t*_ is a *K*-element vector containing the task variables that may affect an animal’s decision on trial *t* ∈ {1, *…, T*}, where **X** = [**x**_1_, *…*, **x**_*T*_]. For the IBL task, **x**_*t*_ could include the contrast values of left and right gratings, as well as stimulus history, a bias term, and other covariates available to the animal during the current trial (Figure 1B). We model the animal’s decision-making process with a Bernoulli generalized linear model (GLM), also known as the logistic regression model. This model characterizes the animal’s strategy on each trial *t* with a set of *K* linear weights **w**_*t*_, where **W** = [**w**_1_, *…*, **w**_*T*_]. Each **w**_*t*_ describes how the different components of the *K*-element input vector **x**_*t*_ affect the animal’s choice on trial *t*. The probability of a rightward decision (*y*_*t*_ = 1) is given by

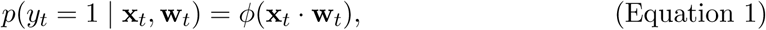

where *ϕ*(·) denotes the logistic function. Unlike standard psychophysical models, which assume that weights are constant across time, we instead assume that the weights evolve gradually over time (Figure 1C). Specifically, we model the weight change after each trial with a Gaussian distribution (Bak et al., 2016; Roy et al., 2018a):

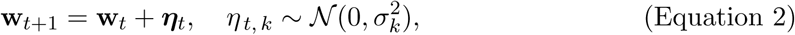

where ***η***_*t*_ is the vector of weight changes on trial *t*, and 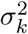 denotes the variance of the changes in the *k*^th^ weight. The rate of change of the *K* different weights in **w**_*t*_ is thus governed by a vector of smoothness hyperparameters *θ* = *{σ*_1_, *…, σ*_*K*_ *}*. The larger the standard deviation *σ*_*k*_, the larger the trial-to-trial changes in the *k*^th^ weight. Note that if *σ*_*k*_ = 0 for all *k*, the weights are constant, and we recover the classic psychophysical model with a fixed set of weights for the entire dataset.

Learning to perform a new task can be formalized under this model by a dynamic trajectory in weight space. Figure 1C-D shows a schematic example of such learning in the context of the IBL task. Here, the hypothetical mouse’s behavior is governed by three weights: a Left Contrast weight, a Right Contrast weight, and a (Choice) Bias weight. The first two weights capture how sensitive the animal’s choice is to left and right gratings, respectively, whereas the bias weight captures a fixed additional bias towards leftward or rightward choices.

In this hypothetical example, the weights evolve over the course of training as the animal learns the task. Initially, during “early training”, the left and right contrast weights are close to zero and the bias weight is large and positive, indicating that the animal pays little attention to the left and right contrasts and exhibits a strong rightward choice bias. As training proceeds, the contrast weights diverge from zero and separate, indicating that the animal learns to compute a difference between right and left contrast. By the “late training” period, left and right contrast weights have grown to equal and opposite values, while the bias weight has shrunk to nearly zero, indicating unbiased, highly accurate performance of the task.

Although we have arbitrarily divided the data into three different periods—designated “early”, “middle” and “late training”—the three weights change gradually after every trial, providing a fully dynamic description of the animal’s decision-making strategy as it evolves during learning. To visualize this strategy, we can compute an “instantaneous psychometric curve” from the weights on any given trial (Figure 1D). These curves describe how the mouse converts the visual stimuli to a probability over choice on any trial, and illustrate the gradual evolution from strongly right-biased toward a high-accuracy strategy in this example. Of course, by incorporating weights on additional task covariates (e.g. choice and reward history), the model can characterize time-varying strategies that are more complex than those captured by the psychometric curve.

### Inferring Weight Trajectories from Data

The goal of our method is to infer the full time-course of an animal’s decision-making strategy from the observed sequence of stimuli and choices over the course of an experiment. To do so, we estimate the animal’s time-varying weights **W** using the dynamic psychophysical model defined above (Equation 1 and Equation 2), where *T* is the total number of trials in the dataset. Each of the *K* rows of **W** represents the trajectory of a single weight across trials, while each column provides the vector of weights governing decision-making on a single trial. Our method therefore involves inferring *K* × *T* weights from only *T* binary decision variables.

To estimate **W** from data, we use a two-step procedure called empirical Bayes (Bishop, 2006). First, we estimate *θ*, the hyperparameters governing the smoothness of the different weight trajectories, by maximizing evidence, which is the probability of choice data given the inputs (with **W** integrated out). Second, we compute the *maximum a posteriori* (MAP) estimate for **W** given the choice data and the estimated hyperparameters 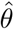. Although this optimization problem is computationally demanding, we have developed fast approximate methods that allow us to model datasets with tens of thousands of trials within minutes on a desktop computer (see Methods for details; see also Figure S1) (Roy et al., 2018a).

To validate the method, we generated a dataset from a simulated observer with *K* = 4 weights that evolved according to a Gaussian random walk over *T* = 5000 trials (Figure 2A). Each random walk had a different standard deviation *σ*_*k*_ (Equation 2), producing weight trajectories with differing average rates of change. We sampled input vectors **x**_*t*_ for each trial from a standard normal distribution, then sampled the observer’s choices **y** according to Equation 1. We applied our method to this simulated dataset, which provided estimates of the 4 hyperparameters 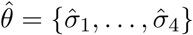 and weight trajectories **Ŵ**. Figure 2A-B shows that our method accurately recovered both the weights **W** and the hyperparameters *θ*.

**Figure 2.**
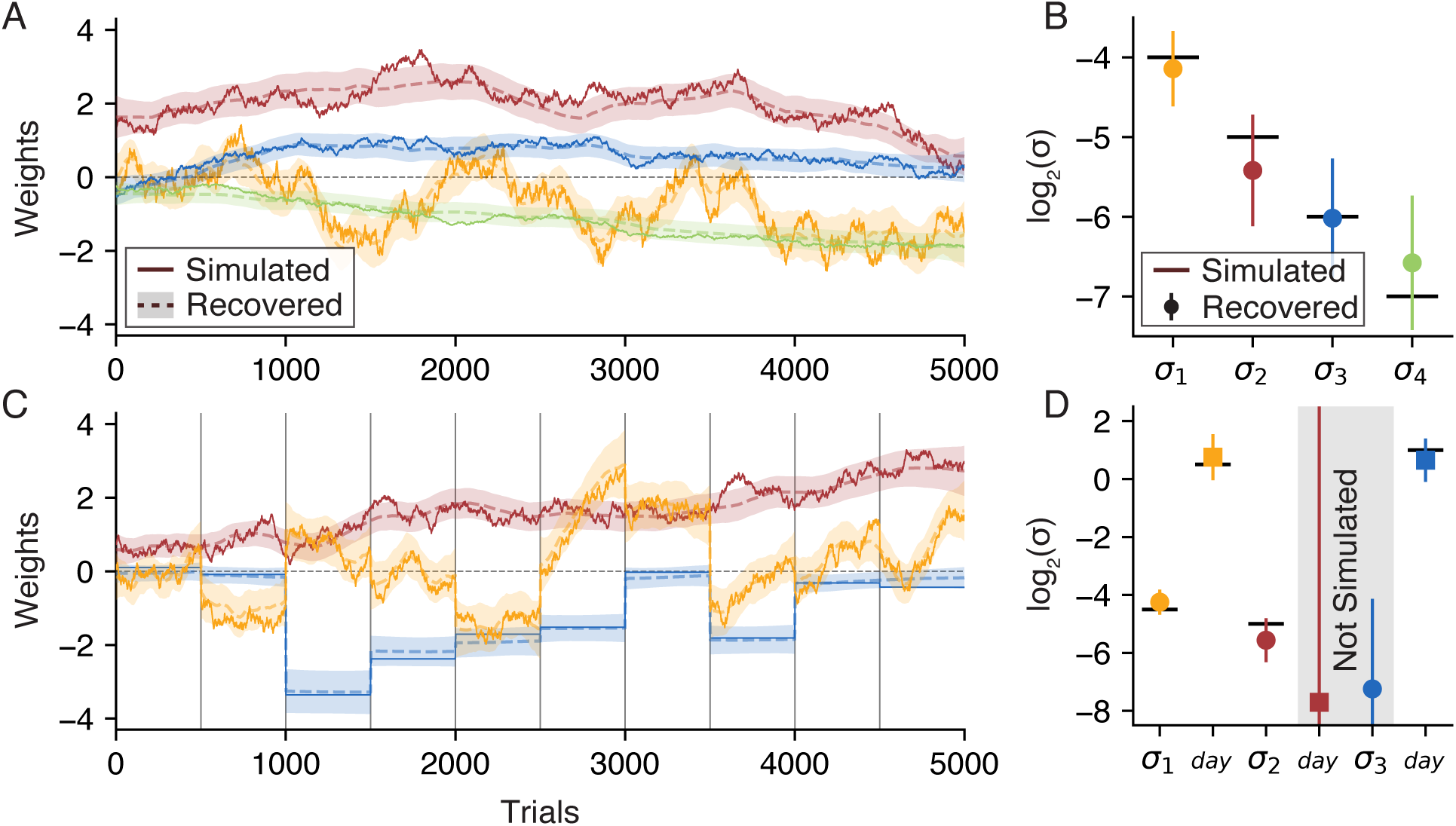
Recovering Psychophysical Weights from Simulated Data. **(A)** To validate our model, we simulated a set of *K* = 4 weights **Ŵ** that evolved for *T* = 5000 trials (solid lines). We\ then use our method to recover weights **W** (dashed lines), surrounded with a shaded 95% credible interval. The full optimization takes less than one minute on a laptop, see Figure S1 for more information. **(B)** In addition to recovering the weights **W** in (A), we also recovered the smoothness hyperparameter *σ*_*k*_ for each weight, also plotted with a 95% credible interval. True values 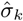 are plotted as solid black lines. **(C)** We again simulated a set of weights as in (A), except now session boundaries have been introduced at 500 trial intervals (vertical black lines), breaking the 5000 trials into 10 sessions. The yellow weight is simulated with a *σ*_day_ hyperparameter much greater than its respective *σ* hyperparameter, allowing the weight to evolve trial-to-trial as well as “jump” at session boundaries. The blue weight has only a *σ*_day_ and no *σ* hyperparameter meaning the weight evolves *only* at session boundaries. The red weight does not have a *σ*_day_ hyperparameter, and so evolves like the weights in (A). **(D)** We recovered the smoothness hyperparameters *θ* for the weights in (C). Though the simulation only has four hyperparameters, the model does not know this and so attempts to recover both a *σ* and a *σ*_day_ hyperparameter for all three weights. The model appropriately assigns low values to the two non-existent hyperparameters (gray shading).

### Augmented Model for Capturing Changes Between Sessions

One limitation of the model described above is that it does not take account of the fact that experiments are typically organized into sessions, each containing tens to hundreds of consecutive trials, with large gaps of time between them. Our basic model makes no allowance for the possibility that weights might change much more between sessions than between other pairs of consecutive trials. For example, if either forgetting or consolidation occurs overnight between sessions, the weights governing the animal’s strategy might change much more dramatically than is typically observed within sessions.

To overcome this limitation, we augmented the model to allow for larger weight changes between sessions. The augmented model has *K* additional hyperparameters, denoted 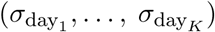, which specify the prior standard deviation over weight changes between sessions or “days”. A large value for 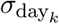 means that the corresponding *k*’th weight can change by a large amount between sessions, regardless of how much it changes between other pairs of consecutive trials. The augmented model thus has 2K hyperparameters, with a pair of hyperparameters 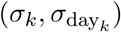 for each of the *K* weights in **w**_*t*_.

We tested the performance of this augmented model using a second simulated dataset which includes session boundaries at 500 trial intervals (Figure 2C). We simulated *K* = 3 weights for *T* = 5000 trials, with the input vector **x**_*t*_ and choices **y**_*t*_ on each trial sampled as in the first dataset. The red weight was simulated like the red weight in Figure 2A, that is, using only the standard *σ* and no *σ*_day_ hyperparameter. Conversely, the blue weight was simulated using *only* a *σ*_day_ hyperparameter, such that the weight is constant within each session, but can “jump” at session boundaries. The yellow weight was simulated with both types of smoothness hyperparameter, allowing it to smoothly evolve within a session as well as evolve more dramatically at session boundaries. Once again, we can see that the recovered weights closely agree with the true weights.

Figure 2D shows the hyperparameters recovered from the second dataset. While the simulated weights had only four hyperparameters (a *σ* for the yellow and red weights, and a *σ*_day_ for the yellow and blue weights), our method inferred both a *σ* and *σ*_day_ hyperparameter for each of the three weights. Thus two of the hyperparameters recovered by the model were not simulated, as indicated by the gray shading. In practice, recovering a value for these unsimulated hyperparameters that is low relative to other hyperparameters will result in an accurate recovery of weights. Thus, the two unsimulated hyperparameters were accurately recovered, as were the four simulated hyperparameters. All subsequent models are fit with *σ*_day_ hyperparameters unless otherwise indicated.

### Characterizing Learning Trajectories in the IBL Task

We now turn to real data, and show how our method can be used to characterize the detailed trajectories of learning in a large cohort of animals. We examined a dataset from the International Brain Lab (IBL) containing behavioral data from *>*100 mice on a standardized sensory decision-making task (see task schematic in Figure 1A) (IBL et al., 2020).

We began by analyzing choice data from the earliest sessions of training. Figure 3A shows the learning curve (defined as the fraction of correct choices per session) for an example mouse over the first several weeks of training. Early training sessions used “easy” stimuli (100% and 50% contrasts) only, with harder stimuli (25%, 12.5%, 6.25%, and 0% contrasts) introduced later as task accuracy improved. To keep the metric consistent, we calculated accuracy only from easy-contrast trials on all sessions.

**Figure 3.**
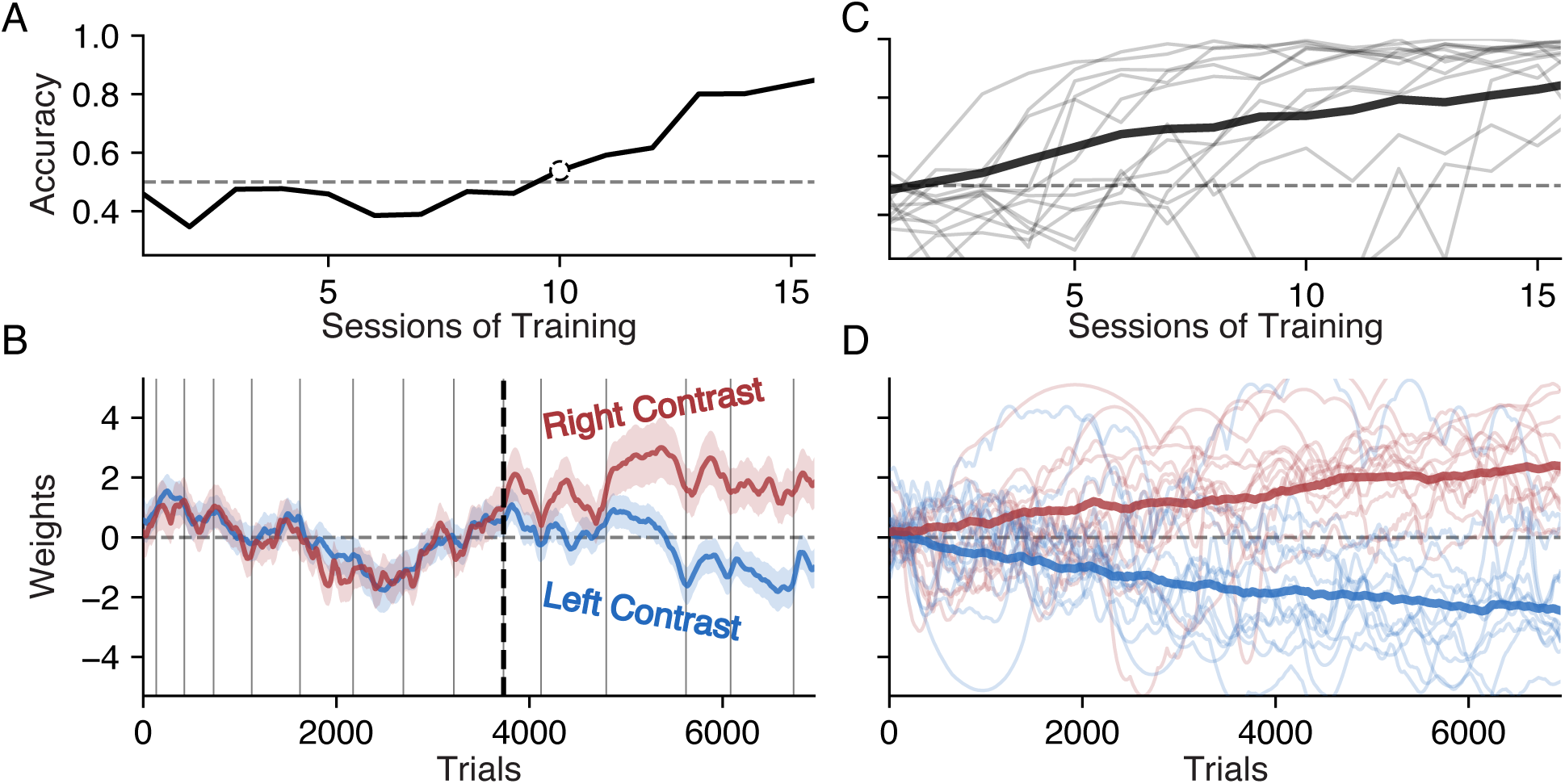
Visualization of Early Learning in IBL Mice. **(A)** The accuracy of an example mouse over the first 16 sessions of training on the IBL task. We calculated accuracy only from “easy” high-contrast (50% and 100%) trials, since lower-contrast stimuli were only introduced later in training. The first session above chance performance (50% accuracy) is marked with a dotted circle. **(B)** Inferred weights for left (blue) and right (red) stimuli governing choice for the same example mouse and sessions shown in (A). Grey vertical lines indicate session boundaries. The black dotted line marks the start of the tenth session, when left and right weights first diverged, corresponding to the first rise in accuracy above chance performance (shown in A). See Figure S2 for models using additional weights. **(C)** Accuracy on easy trials for a random subset of individual IBL mice (gray), as well as the average accuracy of the entire population (black). **(D)** The psychophysical weights for left and right contrasts for the same subset of mice depicted in (C) (light red and blue), as well as the average psychophysical weights of the entire population (dark red and blue) (*σ*_day_ omitted here for visual clarity).

Although traditional analyses stop at coarse performance statistics like accuracy-per-session, the dynamic GLM provides a detailed characterization of the animal’s evolving behavioral strategy at the timescale of single trials. In Figure 3B, we used it to extract time-varying sensory weights on left contrast values (blue) and right-side contrast values (red). These two weights fluctuated together during the first nine sessions, indicating a near-equal probability of making a rightward choice for stimuli on the left and right side of the screen. Positive (negative) fluctuations corresponded to a bias toward rightward (leftward) choices, indicating that the animal’s strategy was not constant across these sessions, even though accuracy remained at chance-level performance. At the start of the tenth session, the two weights began to diverge, corresponding to an increase in accuracy. This separation continued throughout the subsequent six sessions, gradually increasing performance to roughly 80% accuracy by the sixteenth session. The correlated fluctuations of the inferred weights indicate that the animal’s bias changed on a faster timescale than single sessions.

However, the learning trajectory of this example mouse was by no means characteristic of the entire cohort. Figure 3C-D shows the empirical learning curves (above) and inferred weight trajectories (below) from a dozen additional mice selected randomly from the IBL dataset. The light red and blue lines are the right and left contrast weights for individual mice, whereas the dark red and blue lines are the average weights calculated across the entire population. While there is great diversity in the dynamics of the contrast weights of individual mice, we see a smooth and gradual separation of the contrast weights on average.

### Adaptive Bias Modulation in a Non-Stationary Task

Once training has progressed to include contrast levels of all difficulties, the IBL task introduces a final modification to the task. Instead of left and right contrasts appearing randomly with an equal 50:50 probability on each trial, trials are now grouped into alternating “left blocks” and “right blocks”. Within a left block, the ratio of left contrasts to right contrasts is 80:20, whereas the ratio is 20:80 within right blocks. The blocks are of variable length, and sessions sometimes begin with a 50:50 block for calibration purposes.

Using the same example mouse from Figure 3, we extend our view of its task accuracy (on “easy” high-contrast trials) to include the first fifty sessions of training in Figure 4A. The initial gray shaded region indicates training prior to the introduction of bias blocks. The pink outline designates a period of three sessions as “Early Bias Blocks” which includes the last session without blocks and the first two sessions with blocks. The purple outline looks at two arbitrary sessions several weeks of training later, a period designated as “Late Bias Blocks”.

**Figure 4.**
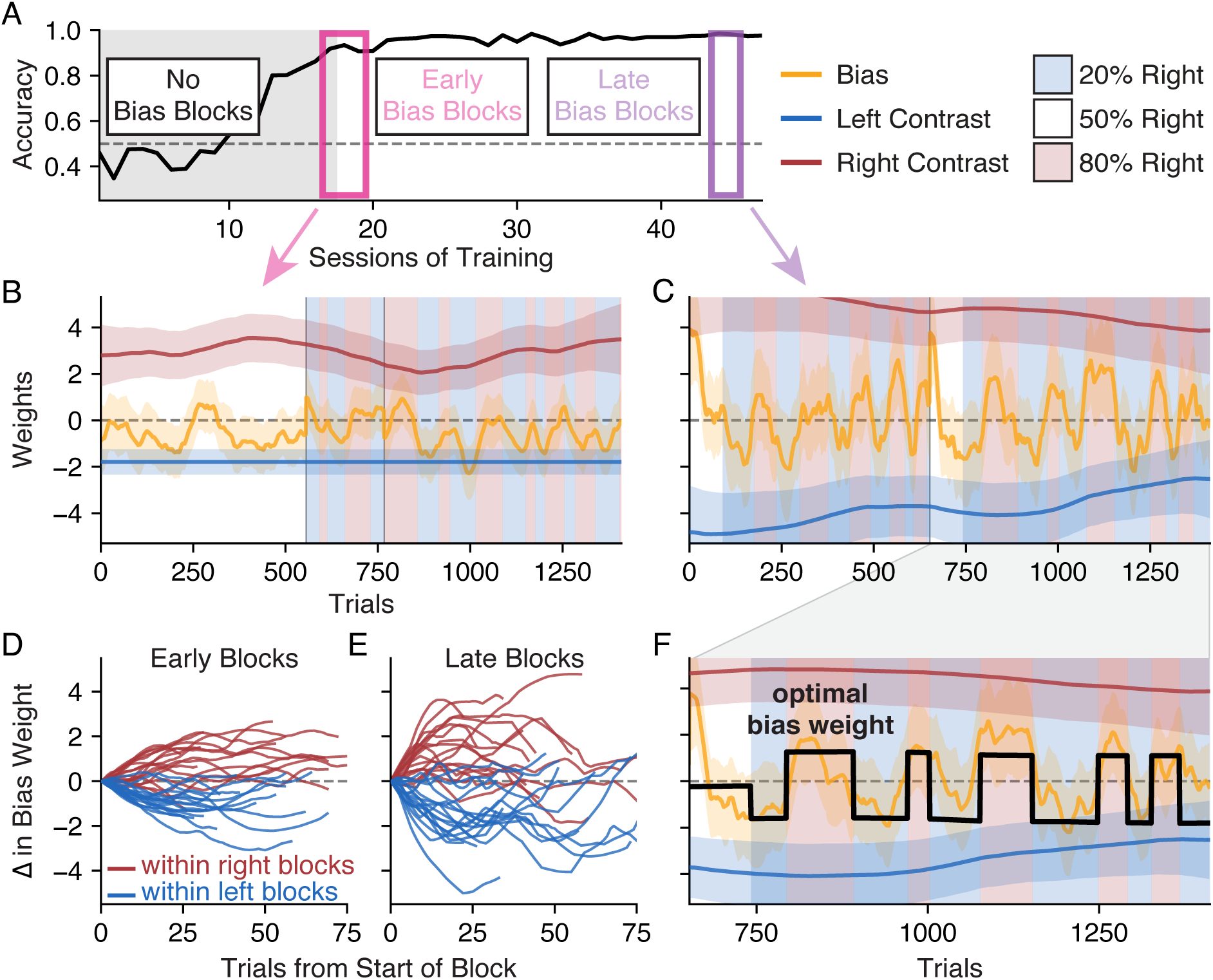
Adaptation to Bias Blocks in an Example IBL Mouse. **(A)** An extension of Figure 3A to include the first months of training (accuracy is calculated only on “easy” high-contrast trials). Starting on session 17, our example mouse was introduced to alternating blocks of 80% right contrast trials (right blocks) and 80% left contrasts (left blocks). The sessions where these bias blocks were first introduced are outlined (pink), as are two sessions in later training where the mouse has adapted to the block structure (purple). **(B)** Three psychophysical weights evolving during the transition to bias blocks, with right (left) blocks in red (blue) shading. Weights correspond to contrasts presented on the left (blue) and right (red), as well as a weight on choice bias (yellow). See Figure S3 for models that parametrize contrast values differently. **(C)** After several weeks of training on the bias blocks, the mouse learns to quickly adapt its behavior to the alternating block structure, as can be seen in the dramatic oscillations of the bias weight in sync with the blocks. (D) The bias weight of our example mouse during the first three sessions of bias block, where the bias weight is chunked by block and each chunk is normalized to start at 0. Even during the initial sessions of bias blocks, the red (blue) lines show that a mild rightward (leftward) bias tended to evolve during right (left) blocks. **(E)** Same as (D) for three sessions during the “Late Bias Blocks” period. Changes in bias weight became more dramatic, tracking stimulus statistics more rapidly, and indicating that the mouse had adapted its behavior to the block structure of the task. **(F)** For the second session from the “Late Bias Blocks” shown in (C), we calculated the *optimal* bias weight (black) given the block transition times and the animal’s sensory stimulus weights. This optimal bias closely matches the empirical bias weight recovered using our model (yellow), indicating that the animal’s strategy was approximately optimal for maximizing reward under the task structure.

In Figure 4B, we apply our method to three “Early Bias Blocks” sessions. Our left and right contrast weights are the same as in Figure 3B (though they now also characterize sensitivity to “hard” as well as “easy” contrast values). We also introduce a third psychophysical weight, in yellow, that tracks sensitivity to choice bias: when this weight is positive (negative), the animal prefers to choose right (left) independent of any other input variable. While task accuracy improves as the right contrast weight grows more positive and the left contrast weight grows more negative, the “optimal” value of the bias weight is naively 0 (no *a priori* preference for either side). However, this is only true when contrasts are presented with a 50:50 ratio and the two contrast weights are of equal and opposite size.

For this mouse, we see that on the last session before the introduction of bias blocks, the bias weight tends to drift around zero and the contrast weights have continued to grow in magnitude from Figure 3B. When the bias blocks commence on the next session, the bias weight does not seem to immediately reflect any change in the stimulus ratio. In Figure 4C, however, we see a clear adaptation of behavior after several weeks of training with the bias block structure. Here the bias weight not only fluctuates more dramatically in magnitude, but is also synchronized with the block structure of the trials.

We examine this phenomenon more precisely in Figure 4D-F, which leverages the results of our method for further analysis. To better examine how our mouse’s choice behavior is changing within a bias block, we can chunk our bias weight according to the start and end of each block. Plotting the resulting chunk (normalized to start at 0) shows how the bias weight changes within a single block. In Figure 4D, we plot all the bias weight chunks from the first three sessions of bias blocks. Chunks of the bias weight that occurred during a left block are colored blue, while chunks occurring during a right block are colored red.

Viewed in this way, we can see that there is some adaptation to the bias blocks even within the first few sessions. Within only a few dozen trials, the mouse’s choice bias tends to slowly drift rightward during right blocks and leftward during left blocks. If we run the same analysis on three sessions near the “Late Bias Blocks” period, we see that this adaptation becomes more dramatic after training (Figure 4E).

We can further analyze the animal’s choice bias in response to the bias blocks by returning to the notion of an “optimal” bias weight. As mentioned before, the naive “optimal” value of a bias weight is zero. However, a non-zero choice bias could *improve* accuracy if, say, the contrast weights are asymmetric (i.e., a left choice bias is useful if the animal is disproportionately sensitive to right contrasts). The introduction of bias blocks further increases the potential benefit of a non-zero bias weight. Suppose that the values of the two contrast weights are so great that the animal can perfectly detect and respond correctly to all contrasts. Even in this scenario, one-in-nine contrasts are 0% contrasts, meaning that (with a bias weight of zero) the animal’s accuracy maxes out at 94.4% (100% accuracy on 8/9 trials, 50% on 1/9). If instead the bias weight adapted perfectly with the bias blocks, the mouse could get the 0% contrasts trials correct with 80% accuracy instead of 50%, increasing its overall accuracy to 97.8%.

Whereas normative models might *derive* a notion of optimal behavior (Tajima et al., 2019), we can use our descriptive model to *calculate* what the “optimal” bias weight would be for each trial and compare it to the bias weight recovered from the data. Here we define an “optimal” bias weight on each trial as the value of the weight that maximizes expected accuracy given that (a the left and right contrasts weights recovered from the data are considered fixed, (b) the precise timings of the block transitions are known, and (c) the distribution of contrast values within each block are known. Under those assumptions, we take the second session of data depicted in Figure 4C and re-plot the psychophysical weights in Figure 4F with the calculated optimal bias weight superimposed in black. Note that the optimal bias weight jumps precisely at each block transition, but also adjusts subtly within a block to account for changes in the contrast weights. We see that, in most blocks, the empirical bias weight (in yellow) matches the optimal bias weight closely. In fact, we can calculate that the mouse would only increase its expected accuracy from 86.1% to 89.3% with the optimal bias weight instead of the empirical bias weight.

### Trial History Dominates Early Behavior in Akrami Rats

To further explore the capabilities of our model, we analyzed behavioral data from another binary decision-making task previously reported in Akrami et al. (2018), where both rats and human subjects were trained on versions of the task (referred to hereafter as “Akrami Rats” and “Akrami Humans”). This auditory parametric working memory task requires a rat to listen to two white noise tones, Tone A then Tone B, of different amplitude and separated by a delay (Figure 5A). If Tone A is louder than Tone B, than the rat must nose poke right to receive a reward, and vice-versa.

**Figure 5.**
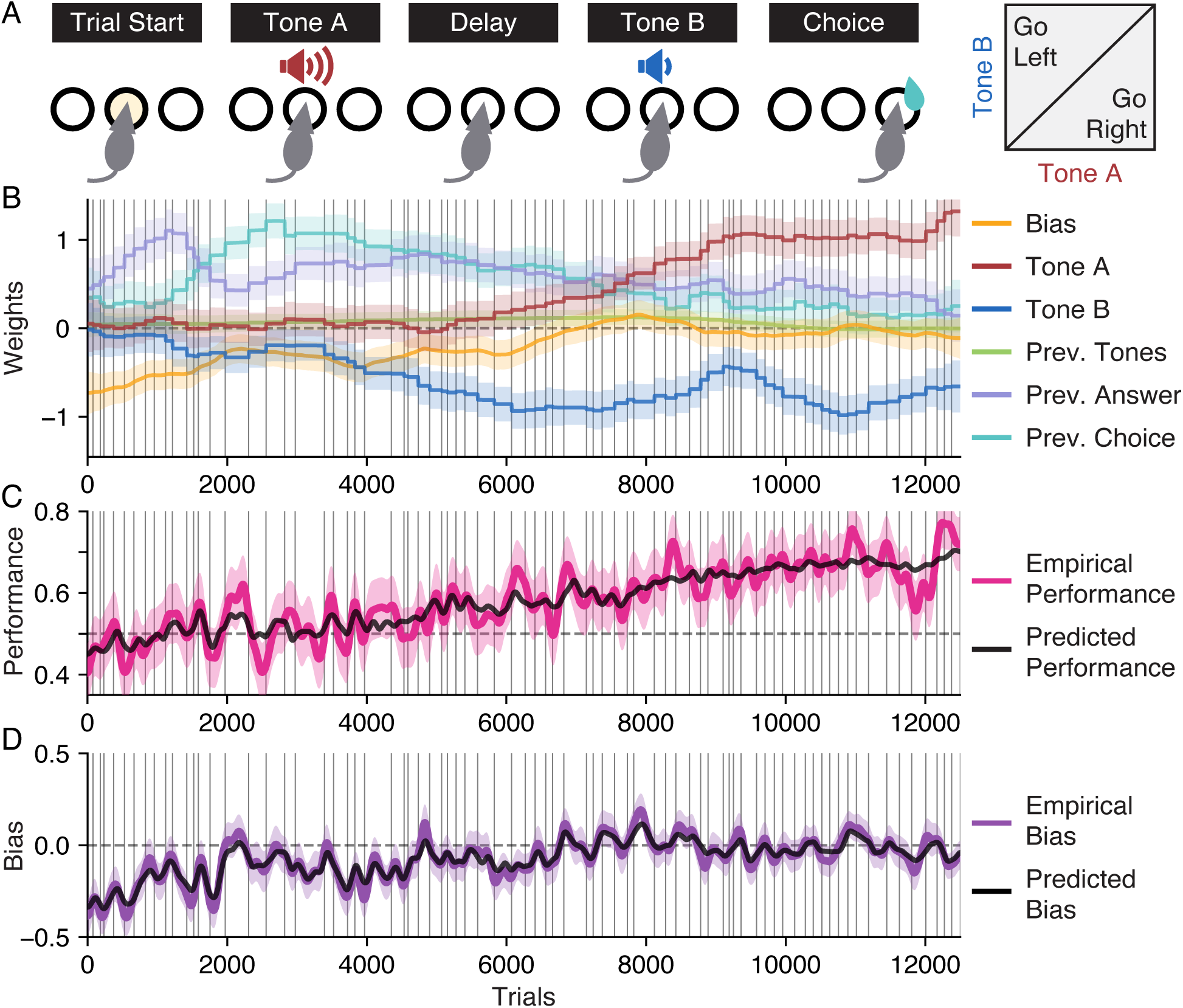
Visualization of Learning in an Example Akrami Rat. **(A)** For this data from Akrami et al. (2018), a delayed response auditory discrimination task was used in which a rat experiences an auditory white noise stimulus of a particular amplitude (Tone A), a delay period, a second stimulus of a different amplitude (Tone B), and finally the choice to go either left or right. If Tone A was louder than Tone B, then a rightward choice triggers a reward, and vice-versa. **(B)** The psychophysical weights recovered from the first 12,500 trials of an example rat. “Prev. Tones” is the average amplitude of Tones A and B presented on the previous trial; “Prev. Answer” is the rewarded (correct) side on the previous trial; “Prev. Choice” is the animal’s choice on the previous trial. Black vertical lines are session boundaries. **(C)** The empirical accuracy of the example rat is plotted in pink, with a 95% confidence interval indicated by the shaded region. We use a cross-validation procedure to calculate the accuracy predicted by the model (black line). Both lines are Gaussian smoothed with *σ* = 50. **(D)** The empirical bias of the example rat is plotted in purple, with a 95% confidence interval indicated by the shaded region. We use a cross-validation procedure to calculate the bias predicted by the model (black line). Both lines are Gaussian smoothed with *σ* = 50. See the Methods for more details about the task, rat subjects, calculation of predicted accuracy and bias, and the cross-validation procedure.

In Figure 5B, we apply our method to the first 12,500 trials of behavior from our example rat. Despite the new task and species, there are several similarities to the results from the IBL mice in Figure 3 and Figure 4. Tones A and B are the task-relevant weights (red and blue, respectively) treated similarly to the left and right contrast variables, while the bias weight (yellow) is the same weight used by the IBL mice. While at most one of the left or right contrast weights was activated on a single trial in the IBL task, however, both the Tone A and B weights are activated on every trial in the Akrami task (the inputs are also parametrized differently, see Methods).

The biggest difference in our application of the method to the Akrami rats is the inclusion of “history regressor” weights; that is, weights which carry information about the previous trial. The Previous Tones variable is the average of Tones A and B from the previous trial (green weight). The Previous (Correct) Answer variable indexes the rewarded side on the previous trial, tracking a categorical variable where {-1,0,+1} = {Left, N/A, Right} (purple weight). The Previous Choice variable indexes the chosen side on the previous trial, also a categorical variable where {-1,0,+1} = {Left, N/A, Right} (cyan weight). History regressor weights are always irrelevant in this task. Despite this, the information of previous trials often influences animal choice behavior, especially early in training (for the impact of history regressors during early learning in an IBL mouse, see Figure S2A).

As can be seen clearly from our example rat, the set of task-irrelevant weights (the three history regressor weights plus the bias weight) dominate behavior early in training. In contrast, the task-relevant weights (on Tones A and B) initialize at zero. However, as training progresses, the task-irrelevant weights shrink while the Tone A and B weights grow to be equal and opposite. Note that the weight on Tone B begins to evolve away from zero very early in training, while the weight on Tone A does not become positive until after tens of sessions. In context of the task, this makes intuitive sense: the association between a louder Tone B and reward on the left is comparatively easy since Tone B occurs immediately prior to the choice. Making the association between a louder Tone A and reward on the right is much more difficult to establish due to the delay period.

Furthermore, the positive value of all three history regressor weights matches our intuitions. The positive value of the Previous (Correct) Answer weight indicates that the animal prefers to go right (left) when the correct answer on the previous trial was also right (left). This is a commonly observed behavior known as a “win-stay/lose-switch” strategy, where an animal will repeat its choice from the previous trial if it was rewarded and will otherwise switch sides. The positive value of the Previous Choice weight indicates that the animal prefers to go right (left) when it also went right (left) on the previous trial. This is known as a “perseverance” behavior: the animal prefers to simply repeat the same choice it made on the previous trial, independent of reward or task stimuli. The slight positive value of the Previous Tones weight indicates that the animal is biased toward the right when the tones on the previous trial were louder than average, the same effect produced by Tone A. This corroborates an important finding from the original paper: the biases seen in choice behavior are consistent with the mouse’s memory of Tone A being corrupted by recent sensory history (Akrami et al. (2018), though note that the analysis there was done on post-training behavior and uses the 20-50 most recent trials to calculate an average previous tones term; see also Papadimitriou et al. (2015); Lu et al. (1992)).

While our simulated datasets in Figure 2 validated our method’s ability to recover weights from data generated by our own model, we would like to confirm that the weights recovered by our method accurately characterize behavior in real data. To accomplish this, we use the weights recovered from our model to predict the values of more conventional behavioral metrics. Figure 5C shows the empirical accuracy of our example rat (pink) with a 95% confidence interval (shaded pink). Combining the task variable with our weights from Figure 5B according to Equation 1, we can calculate *P* (Correct Answer) for each trial to infer the model’s predicted accuracy. This predicted accuracy is overlaid in black in Figure 5C, and we see that our model’s prediction largely agrees with the empirical measure.

This validation of the model is repeated in Figure 5D, here calculating empirical and predicted choice bias. Again we see a close agreement between the empirical predicted measure. Note that this measure of bias is distinct from the specific bias weight (Figure 5B, yellow), as it is calculated using all the weights of the model. When only taking into consideration the task stimuli and the animal’s choice, there are many strategies that can masquerade as a choice bias. For example, a “perseverance” behavior where the animal repeats it’s choice from the previous trial could also look like a preference for a particular side in the short-term. The bias weight in our model captures choice bias *independent* of the other behaviors.

In calculating the predicted accuracy and bias, it is not appropriate to evaluate on data the model used for optimization. To avoid this, we use a 10-fold cross-validation procedure where the model is fit using only 90% of trials and predictive metrics are calculated using the remaining 10% of trials. For more details regarding this cross-validation procedure as well as the precise calculation of accuracy and bias, see the Methods.

### Behavioral Trends Are Shared Across the Population of Akrami Rats

By applying our method to the entire population of Akrami rats, we can uncover trends and points of variation in training. Figure 6 looks at the same set of weights used in Figure 5B. Here, trajectories are calculated for the first 20,000 trials of training for each rat. Each plot consists of the individual weight trajectories from the whole population of rats as well as the average trajectory. Figure 6A shows the weights for both Tone A and Tone B. The observation from our example rat that Tone B grows before the Tone A weight appears to hold uniformly across the population. There is more extensive variation in the bias weights of the population as seen in Figure 6B, though the variation is greatest early in training and at all points averages out to approximately 0. The slight positive value of the Previous Tones weight is highly consistent across all rats and constant across training, seen in Figure 6C. The prevalence of both “win-stay/lose-switch” and “perseverance” behaviors across the population can be clearly seen in the positive values of the Previous (Correct) Answer and Choice weights in Figure 6D and E, though there is substantial variation in the dynamics of these history regressors.

**Figure 6.**
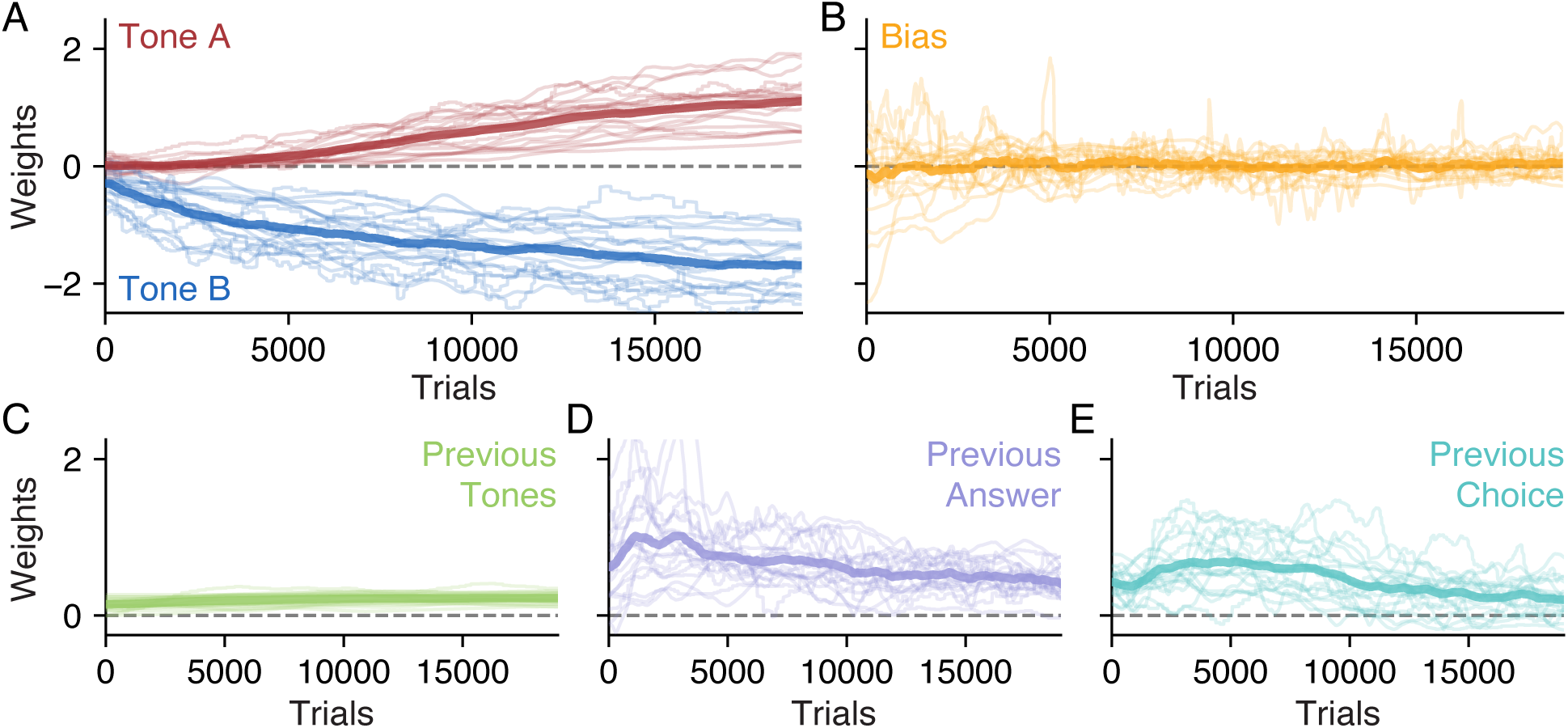
Population Psychophysical Weights from Akrami Rats. The psychophysical weights during the first *T* = 20000 trials of training, plotted for all rats in the population (light lines), plus the average weight (dark line): **(A)** Tone A and Tone B, **(B)** Bias, **(C)** Previous Tones, **(D)** Previous (Correct) Answer, and **(E)** Previous Choice.

### In Contrast to Rats, Human Behavior is Stable

In addition to training rats on a sensory discrimination task, Akrami et al. (2018) also adapted the same task for human subjects. The modified task also requires a human subject to discriminate two tones, though the human chooses with a button instead of a nose-poke and is rewarded with money (Figure 7A). The weights from an example human subject are shown in Figure 7B, whereas the weights from all the human subjects are shown together in Figure 7C.

**Figure 7.**
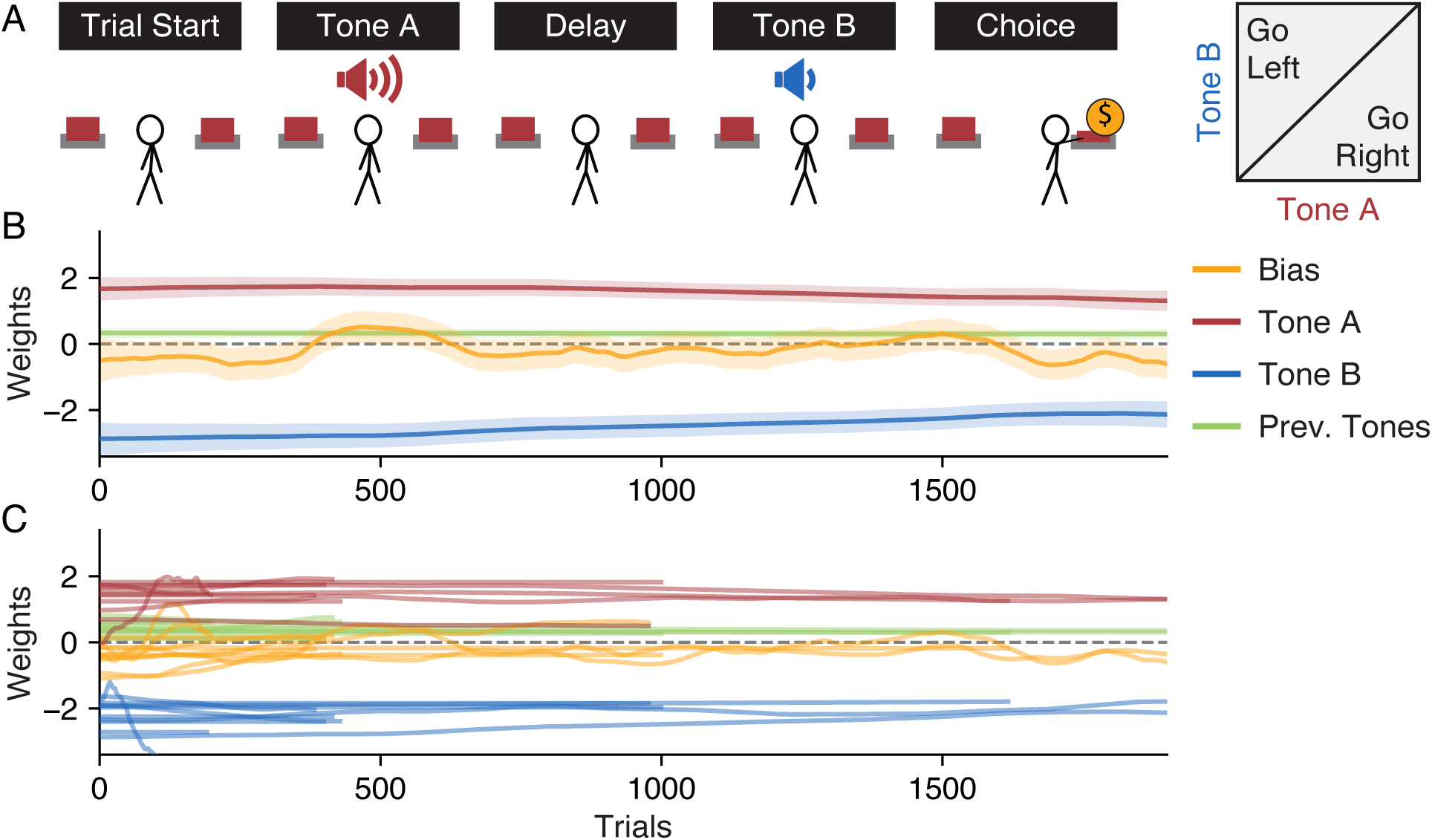
Population Psychophysical Weights from Akrami Human Subjects. **(A)** The same task used by the Akrami rats in Figure 5A, adapted for human subjects. **(B)** The weights for an example human subject. Human behavior is not sensitive to the previous correct answer or previous choice, so corresponding weights are not included in the model (see Figure S4 for a model which includes these weights). **(C)** The weights for the entire population of human subjects. Human behavior was evaluated in a single session of variable length.

It is useful to contrast the weights recovered from the Akrami human subjects to the weights recovered from the Akrami rats. Since the rules of the task are explained to human subjects, one would intuitively expect that human weights would initialize at “correct” values corresponding to high performance and would remain constant throughout training. These, however, are assumptions. An advantage of our method is that no such assumptions need to be made: the initial values and stability of the weights is determined entirely from the data. From Figure 7C, we see that the model does indeed confirm our intuitions without the need for us to impose them *a priori*. All four weights remain relatively stable throughout training, though choice bias does oscillate near zero for some subjects. Two of the history regressor weights that dominated early behavior in rats, Previous (Correct) Answer and Choice, are not used by humans in the population and so are omitted here (see Figure S4). The slight positive weight on Previous Tones remains for all subjects, however, and there is a slight asymmetry in the magnitudes of the Tone A and Tone B weights in many subjects.

### Including History Regressors Boosts Predictive Power

Just as we were able to leverage the results of our analysis of the IBL mice to examine the impact of the bias block structure on the choice bias of an example mouse in Figure 4, we can also extend our analysis of the Akrami rats to quantify the importance of history regressors in characterizing behavior. Using our example rat from Figure 5, we wish to quantify the difference between a model that includes the three history regressors and another model without them (one with weights for only Tone A, Tone B, and Bias). To do this, we calculate the model’s predicted accuracy at each trial. The predicted accuracy on a trial *t* can be defined as max(*P* (Right), *P* (Left)), calculated using Equation 1. Note that for the predictions to be valid, we determine predicted accuracy using weights produced from a cross-validation procedure (Methods).

In Figure 8A, we refit a model to the same data shown in Figure 5B using only Bias, Tone A, and Tone B weights. Next, we bin the trials according to the model’s predicted accuracy, as defined above. Finally, for the trials within each bin, we plot the average predicted accuracy on the x-axis and the empirical accuracy of the model (the fraction of trials in a bin where the prediction of the model matched the choice the animal made) on the y-axis. Points that lie below the dotted identity line represent over-confident predictions from the model, whereas points above the line indicate under-confidence. The fact that points (shown with 95% confidence intervals on their empirical accuracy) lie along the dotted line further validates the accuracy of the model.

**Figure 8.**
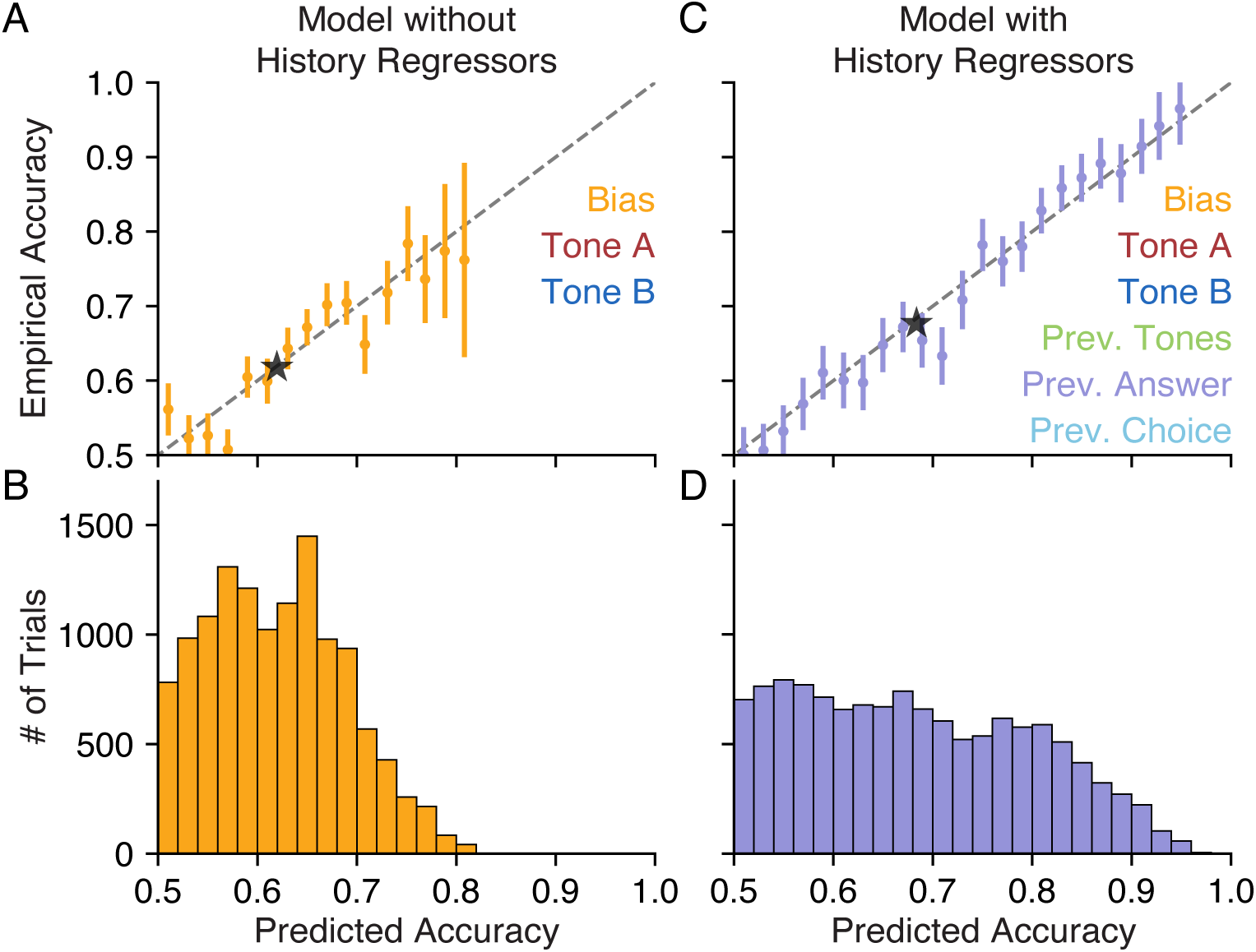
History Regressors Improve Model Accuracy for an Example Akrami Rat. **(A)** Using a model of our example rat that omits history regressors, we plot the empirical accuracy of the model’s choice predictions against the model’s cross-validated predicted accuracy. The black dashed line is the identity, where the predicted accuracy of the model exactly matches the empirical accuracy (i.e., points below the line are *overconfident* predictions). The animal’s choice is predicted with 61.9% confidence on the average trial, precisely matching the model’s empirical accuracy of 61.9% (black star). Each point represents data from the corresponding bin of trials seen in (B). Empirical accuracy is plotted with a 95% confidence interval. See Methods for more information on the cross-validation procedure. **(B)** A histogram of trials binned according to the model’s predicted accuracy. **(C)** Same as (A) but for a model that also includes three additional weights on history regressors: Previous Tones, Previous (Correct) Answer, and Previous Choice. We see that data for this model extends into regions of higher predicted and empirical accuracy, as the inclusion of history regressors allows the model to make stronger predictions. The animal’s choice is predicted with 68.4% confidence on the average trial, slightly overshooting the model’s empirically accuracy of 67.6% (black star). **(D)** Same as (B), but for the model including history regressors.

The histogram in Figure 8B shows the number of trials in each of the bins used in Figure 8A. We can see that almost no choices are predicted with greater than 80% certainty. The black star in Figure 8A shows the average predictive accuracy for this model (61.9%) and corresponding empirical accuracy (also 61.9%).

Adding back the three history regressor weights, we first check if our model is still making valid predictions. Figure 8C shows that most of our data points still lie along the dotted identity line, meaning that the model’s predictions remain well-calibrated. The data points also extend further along the diagonal—accounting for the impact of the previous trial on choice behavior allows the model to make stronger predictions. Examining Figure 8D, we see that a significant fraction of trials now have a predicted accuracy greater than the strongest predictions made by the model without history regressors. In fact, choices on some trials can be predicted with near certainty, with over 95% confidence. As the black star in Figure 8C indicates, the inclusion of history regressors improves the predicted accuracy of the model to 68.4% (slightly overconfident relative to the average empirical accuracy of 67.6%).

## Discussion

Training experimental subjects to achieve stable, interpretable behavior is the practical first step in addressing many neuroscientific questions. The motivation is clear: to make sense of neural activity, we must first make sense of the behavior that activity produces (Carandini, 2012). Despite the current excitement surrounding new neural analysis techniques (Williams et al., 2020; Duncker and Sahani, 2018; Meshulam et al., 2019; Semedo et al., 2019; Kobak et al., 2016; Gao et al., 2016), however, analysis of the behavior facilitating those techniques has received comparatively little attention. The standard suite of behavioral analysis tools is ill-equipped to capture dynamic and complex behavioral strategies. To address this burgeoning research bottleneck, we have presented a model of time-varying psychophysical weights that captures the evolution of complex decision-making strategies. Across two tasks and three species, we demonstrated that our method not only characterizes complex behavior at a trial-by-trial resolution, but also serves as a foundation for more targeted analyses.

The tools typically used to quantify decision-making behavior, such as learning curves (Figure 3A) and psychometric functions (Figure 1D), provide a limited and static view of behavior. Decision-making is a dynamic process that often entails the use of many different behavioral strategies to achieve task proficiency. These dynamics are especially important in a time-varying task environment, often with a continual learning to account for a potentially dynamic environment (Piet et al., 2018). Research geared to study the learning process in the context of neural recordings has received steady attention (Schultz et al., 1997; Dayan and Balleine, 2002; O’Doherty et al., 2003; Daw and Doya, 2006). On the other hand, general methods for characterizing the learning process of animals purely from choice behavior are less common, despite their broad applicability and potential practical applications. Our work highlights the richness of information that can be inferred from the trial-to-trial dynamics of decision-making behavior.

Our method provides a highly informative, as well as efficient, way of characterizing behavioral dynamics, highlighting the richness of information that can be inferred from the trial-to-trial dynamics of decision-making behavior. Our work offers significant improvement from previous methods. Work from Smith et al. (2004) used choice behavior to estimate learning curves, but stopped short of linking changes in performance to specific behavioral strategies. Past work from Frund et al. (2014) introduced an explicit model of inter-trial dependence in decision-making, but was limited to the paradigm of assuming a single, true psychometric function. Our approach builds upon a state-space approach for dynamic tracking of behavior (Bak et al., 2016), utilizing the decoupled Laplace method (Wu et al., 2017) to scale up analysis and make it practical for modern behavioral datasets (Methods). In particular, the efficiency of our algorithm allows for routine analysis of large behavior datasets, with tens of thousands of trials, within minutes on a laptop (see Figure S1).

The breadth of opportunities indicated by the range of analyses presented here leads us to anticipate several use cases for our model. First, experimenters training animals on binary decision-making tasks can use the model to better understand the diverse range of behavioral strategies seen in early training. This should facilitate the design and validation of new training strategies, which could ultimately open the door to more complex tasks and faster training times. Second, investigations into the learning process itself are made more accessible as datasets of training data, typically left unanalyzed, can now be easily explored and mined for insight into the learning process. Third, the psychophysical weights of our model lend themselves easily to downstream analyses. Our general method can enable more targeted investigations, acting as one step in a larger analysis pipeline (as in Figure 4 and Figure 8). To facilitate these uses, we have released our method as PsyTrack, a publicly available Python package (Roy et al., 2018b). The Google Colab notebook accompanying this work provides many flexible examples and can serve as a template for adaptation to new datasets (Methods).

The two assumptions of our model, that (i) decision-making behavior can be described by a set of GLM weights, and (ii) that these weights evolve smoothly over training, are well-validated in the datasets explored here. However, these assumptions may not be true for all datasets. Behaviors which change suddenly may not be well described by the smoothly evolving weights in our model, though allowing for weights to evolve more dramatically between sessions can mitigate this model mismatch. Determining which input variables to include and how they ought to be parameterized can also be challenging. For example, task-irrelevant covariates of decision-making typically include the history of previous trials (Akrami et al., 2018; Frund et al., 2014; Corrado et al., 2005), which is not always clearly defined; depending on the task, the task-relevant feature may also be a pattern of multiple stimulus units (Murphy et al., 2008). However, this flexibility gives our model the potential to account for a wide variety of behavioral strategies. In the face of these potential pitfalls, it is important to validate the results of our model. Thus, we have provided comparisons to more conventional measures of behavior to help assess the accuracy of our model (see Figure 5 and Figure 8). Furthermore, the Bayesian setting of our modeling approach provides posterior credible intervals for both weights and hyperparameters, allowing for uncertainty in our inferences about behavior to be quantified.

The ability to quantify complex and dynamic behavior at a trial-by-trial resolution enables exciting future opportunities for animal training. As suggested in Bak et al. (2016), the descriptive model of behavior we build here could be extended to an explicit model of learning that makes predictions as to how behavior would change in response to particular stimuli and choices. Ultimately, this could guide the creation of automated optimal training paradigms that can present the specific stimuli to maximize learning. There are also opportunities to extend the application of the method beyond binary decision-making tasks to the multinomial setting where additional choices (or non-choices, e.g. error trials) could also be included in the model (Churchland et al., 2008; Bak and Pillow, 2018). Our work opens up the path toward a more rigorous understanding of the behavioral dynamics at play as animals learn. As researchers continue to ask challenging questions, new animal training tasks will grow in number and complexity. We expect our method to guide those looking to better understand the dynamic behavior of their experimental subjects.

## ACKNOWLEDGEMENTS

The authors thank A. Churchland, A. Pouget, M. Carandini, and A. Urai for feedback on the manuscript. We also thank K. Osorio and J. Teran for animal and laboratory support in collecting the rat high-throughput data. This work was supported by grants from: (IBL) – the Simons Collaboration on the Global Brain and the Wellcome Trust; (NAR & JWP) – the Simons Collaboration on the Global Brain (SCGB AWD543027), the NIH BRAIN initiative (NS104899,R01EB026946), and a U19 NIH-NINDS BRAIN Initiative Award (5U19NS104648).

## AUTHOR CONTRIBUTIONS

Conceptualization, N.A.R., J.H.B., J.W.P.; Methodology, N.A.R., J.H.B., J.W.P.; Software, N.A.R.; Formal Analysis, N.A.R.; Investigation, N.A.R., I.B.L., A.A.; Resources, I.B.L., A.A., C.D.B., J.W.P.; Data Curation, N.A.R., I.B.L., A.A.; Writing – Original Draft, N.A.R.; Writing – Review & Editing, N.A.R., J.H.B., I.B.L., A.A., C.D.B., J.W.P.; Visualization, N.A.R.; Supervision, J.W.P.; Project Administration, N.A.R., J.W.P.; Funding Acquisition, I.B.L., J.W.P.

## METHODS

Detailed methods are provided in the online version of this paper and include the following:

### KEY RESOURCES TABLE

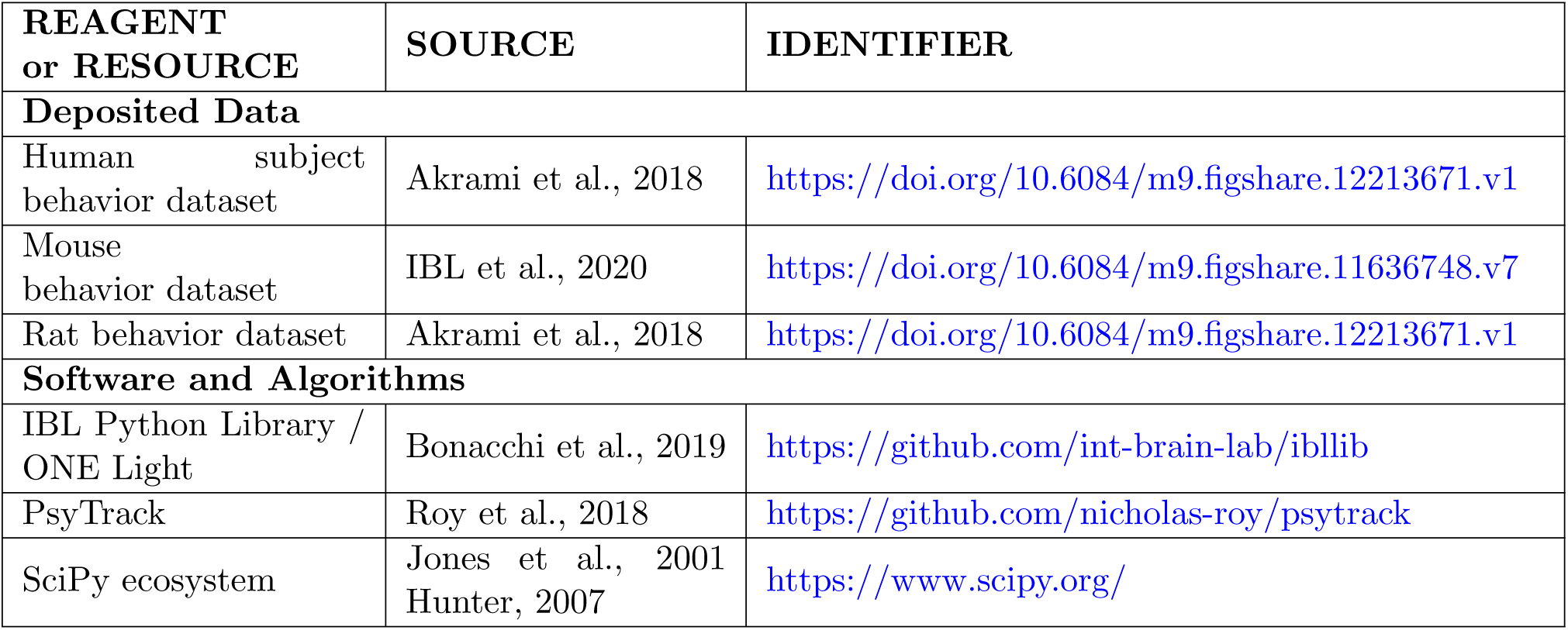

| REAGENT<br>or RESOURCE | SOURCE | IDENTIFIER |
| --- | --- | --- |
| **Deposited Data** |  |  |
| Human subject<br>behavior dataset | Akrami et al., 2018 | https://doi.org/10.6084/m9.figshare.12213671.v1 |
| Mouse<br>behavior dataset | IBL et al., 2020 | https://doi.org/10.6084/m9.figshare.11636748.v7 |
| Rat behavior dataset | Akrami et al., 2018 | https://doi.org/10.6084/m9.figshare.12213671.v1 |
| **Software and Algorithms** |  |  |
| IBL Python Library /<br>ONE Light | Bonacchi et al., 2019 | https://github.com/int-brain-lab/ibllib |
| PsyTrack | Roy et al., 2018 | https://github.com/nicholas-roy/psytrack |
| SciPy ecosystem | Jones et al., 2001<br>Hunter, 2007 | https://www.scipy.org/ |

### LEAD CONTACT AND MATERIALS AVAILABILITY

Further information and requests for resources should be directed to and will be fulfilled by the Lead Contact, Nicholas A. Roy.

### EXPERIMENTAL MODEL AND SUBJECT DETAILS

#### Mouse Subjects

101 experimental subjects were C57BL/6J mice obtained from Jackson Laboratory or Charles River. All procedures and experiments were carried out in accordance with the local laws and following approval by the relevant institutions such as the Animal Welfare Ethical Review Body in the UK and the Institutional Animal Care and Use Committee in the US. This data was first reported in IBL et al. (2020).

#### Rat Subjects

19 experimental subjects were Long–Evans rats (*Rattus norvegicus*) between the ages of 6 and 24 months. Animal use procedures were approved by the Princeton University Institutional Animal Care and Use Committee and carried out in accordance with National Institutes of Health standards. This data was first reported in Akrami et al. (2018).

#### Human Subjects

11 human subjects (8 males and 3 females, aged 22–40) were tested and all gave their informed consent. Participants were paid to be part of the study. The consent procedure and the rest of the protocol were approved by the Princeton University Institutional Review Board. This data was first reported in Akrami et al. (2018).

### METHOD DETAILS

#### Optimization: Psychophysical Weights

Our method requires that weight trajectories be inferred from the response data collected over the course of an experiment. This amounts to a very high-dimensional optimization problem when we consider models with several weights and datasets with tens of thousands of trials. Moreover, we wish to learn the smoothness hyperparameters *θ* = *{σ*_1_, *…, σ*_*K*_} in order to determine how quickly each weight evolves across trials. The theoretical framework of our approach was first introduced in Bak et al. (2016). The statistical innovations facilitating application to large datasets, as well as the initial release of our Python implementation PsyTrack (Roy et al., 2018b), were first presented in Roy et al. (2018a).

We describe our full inference procedure in two steps. The first is optimizing for a weight trajectories **W** given a fixed set of hyperparameters, while the second step optimizes for the hyperparameters *θ* given a fixed set of weights. The full procedure involves alternating between the two steps until both weights and hyperparameters converge.

For now, let **W** denote the massive weight vector formed by concatenating all of the *K* individual length-*T* trajectory vectors, where *T* is the total number of trials. We then define ***η*** = *D***w**, where *D* is a block-diagonal matrix of *K* identical *T* × *T* difference matrices (i.e., 1 on the diagonal and −1 on the lower off-diagonal), such that ***η***_*t*_ = **w**_*t*_ − **w**_*t*−1_ for each trial *t*. Because the prior on ***η*** is simply *𝒯* (**0**, Σ), where Σ has each of the 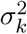 stacked *T* times along the diagonal, the prior for **w** is *𝒯* (**0**, *C*) with *C*^−1^ = *D*^T^Σ^−1^*D*. The log-likelihood is simply a sum of the log probability of the animal’s choice on each trial, 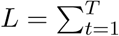 log *p*(*y*_*t*_|**x**_*t*_, **w**_*t*_).

The log-posterior is then given by

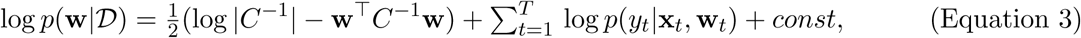

where 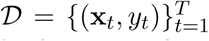 is the set of user-defined input features (including the stimuli) and the animal’s choices, and *const* is independent of **w**.

Our goal is to find the **w** that maximizes this log-posterior; we refer to this *maximum a posteriori* (MAP) vector as **w**_MAP_. With *TK* total parameters (potentially 100’s of thousands) in **w**, most procedures that perform a global optimization of all parameters at once (as done in Bak et al. (2016)) are not feasible; for example, related work has calculated trajectories by maximizing the likelihood using local approximations (Smith et al., 2004). Whereas the use of the Hessian matrix for second-order methods often provides dramatic speed-ups, a Hessian of (*TK*)^2^ parameters is usually too large to fit in memory (let alone invert) for *T >* 1000 trials.

On the other hand, we observe that the Hessian of our log-posterior is sparse:

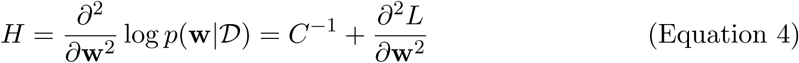

where *C*^−1^ is a sparse (banded) matrix, and *∂*^2^*L/∂***w**^2^ is a block-diagonal matrix. The block diagonal structure arises because the log-likelihood is additive over trials, and weights at one trial *t* do not affect the log-likelihood component from another trial *t*^′^.

We take advantage of this sparsity, using a variant of conjugate gradient optimization that only requires a function for computing the product of the Hessian matrix with an arbitrary vector (Nocedal and Wright, 2006). Since we can compute such a product using only sparse terms and sparse operations, we can utilize quasi-Newton optimization methods in SciPy to find a global optimum for **w**_MAP_, even for very large *T* (Jones et al., 2001).

##### Algorithm 1 Optimizing hyperparameters with the decoupled Laplace approximation

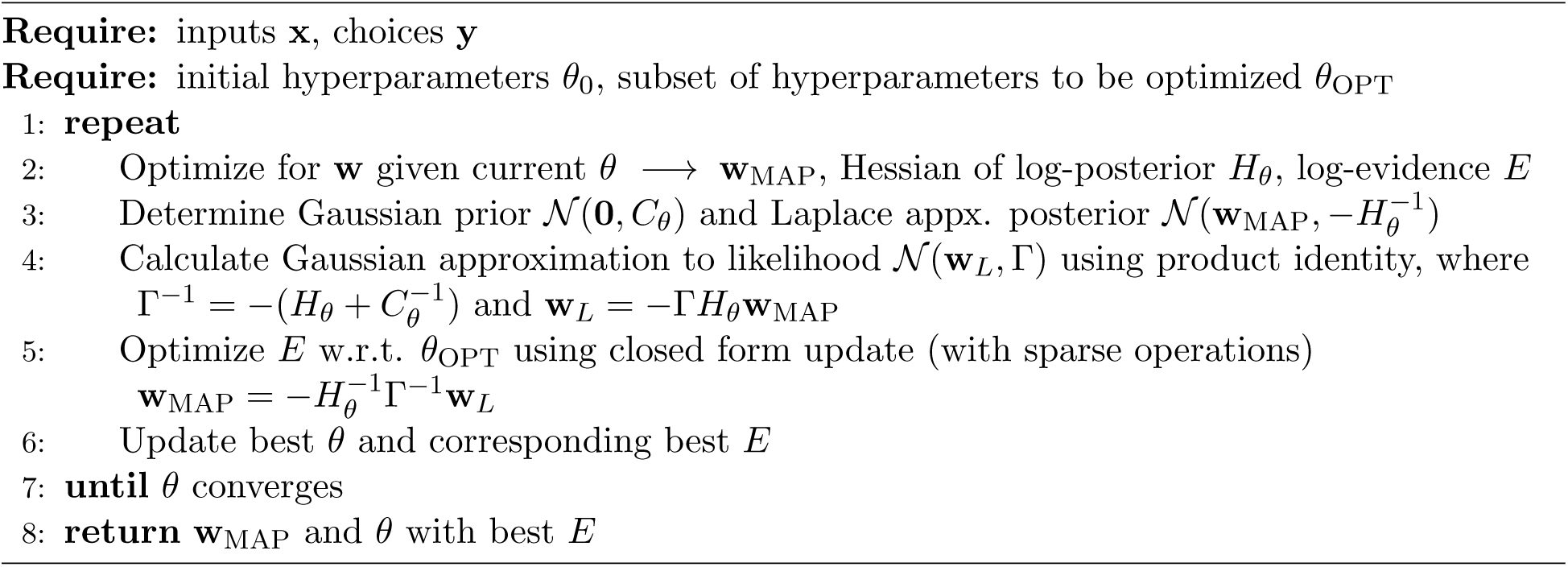

#### Optimization: Smoothness Hyperparameters

So far we have addressed the problem of finding a global optimum for **w, w**_MAP_, given a specific hyperparameter setting *θ*; now we must also find the optimal hyperparameters. A common approach for selecting hyperparameters would be to optimize for cross-validated log-likelihood. Given the potential number of different smoothness hyperparameters and the computational expense of calculating **w**_MAP_, this is not feasible. We turn instead to an optimization of the (approximate) marginal likelihood, or model evidence, called empirical Bayes (Bishop, 2006).

To select between models optimized with different *θ*, we use a Laplace approximation to the posterior, *p*(**w**|*𝒟, θ*) ≈ *𝒩* (**w**|**w**_MAP_, −*H*^−1^), to approximate the marginal likelihood as in (Sahani and Linden, 2003):

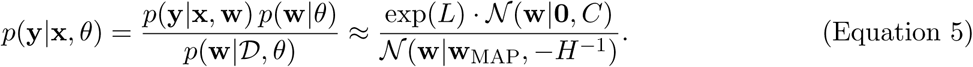

Naive optimization of *θ* requires a re-optimization of **w** for every change in *θ*, strongly restricting the dimensionality of tractable *θ*. Under such a constraint, the simplest approach is to reduce all *σ*_*k*_ to a single *σ*, thus assuming that all weights have the same smoothness (as done in Bak et al. (2016)).

Here we use the decoupled Laplace method (Wu et al., 2017) to avoid the need to re-optimize for our weight parameters after every update to our hyperparameters by making a Gaussian approximation to the likelihood of our model. This optimization is given in Algorithm 1. By circumventing the nested optimizations of *θ* and **w**, we can consider larger sets of hyperparameters and more complex priors over our weights (e.g. *σ*_day_) within minutes on a laptop (see Figure S1). In practice, we also parametrize *θ* by fixing *σ*_*k,t*=0_ = 16, an arbitrary large value that allows the likelihood to determine **w**_0_ rather than forcing the weights to initialize near some predetermined value via the prior.

#### Selection of Input Variables

The variables that make up the model input **x** are entirely user-defined. The decision as to what variables to include when modeling a particular dataset can be determined using the approximate log-evidence (log of Equation 5). The model with the highest approximate log-evidence would be considered best, though this comparison could also be swapped for a more expensive comparison of cross-validated log-likelihood (using the cross-validation procedure discussed below).

Non-identifiability is another issue that should be taken into account when selecting the variables in the model. A non-identifiability in the model occurs if one variable in **x** is a linear combination of some subset of other variables, in which case there are infinite weight values that all correspond to a single identical model. Fortunately, the posterior credible intervals on the weights will help indicate that a model is in a non-identifiable regime — since the weights can take a wide range of values to represent the same model, the credible intervals will be extremely large on the weights contributing to the non-identifiability. See Figure S2B for an example and further explanation.

#### Parameterization of Input Variables

It is important that the variables used in **x** are standardized such that the magnitudes of different weights are comparable. For categorical variables, we constrain values to be {−1, 0, +1}. For example, the Previous Choice variable is coded as a −1 if the choice on the previous trial was left, +1 if right, and 0 if there was no choice on the previous trial (e.g. on the first trial of a session). Additionally, variables depending on the previous trial can be set to 0 if the previous trial was a mistrial. Mistrials (instances where the animal did not complete a trial, e.g., by violating a “go” cue) are otherwise omitted from the analysis. The choice bias is fixed to be a constant +1.

Continuous variables can be more difficult to parameterize appropriately. In the Akrami task, each of the variables for Tone A, Tone B, and Previous Tones are standardized such that the mean is 0 and the standard deviation is 1. The left and right contrast values used in the IBL task present a more difficult normalization problem. Suppose the contrast values were used directly. This would imply that a mouse should be twice as sensitive to a 100% contrast than to a 50% contrast. Empirically, however, this is not the case: mice tend to have little difficulty distinguishing either and perform comparably on these “easy” contrast levels. Nonetheless, both contrast values are used as input to the same contrast weight and so the model will always predict a significant difference in behavior between 50% and 100% contrasts.

Rather than use the raw contrast values as input, our model can be improved by reparameterizing contrasts to better reflect the *perceived* relative differences between contrast values. For example, Busse et al. (2011) used similar visual contrasts to investigate the contrast response function from activity in the visual cortex of anesthetized mice. In this paper, we use a constant transformation of the contrast values for all mice at all points in training (though this could be tuned for each model to maximize the log-evidence). The following tanh transformation of the contrasts *x* has a free parameter *p* which we set as *p* = 5 throughout the paper: 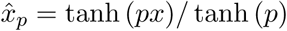. Specifically, this maps the contrast values from [0, 0.0625, 0.125, 0.25, 0.5, 1] to [0, 0.302, 0.555, 0.848, 0.987, 1]. See Figure S3 for a worked example of why this parametrization is necessary.

#### IBL Task

Here we review the relevant features of the task and mouse training protocol from the International Brain Lab task (IBL task). Please refer to IBL et al. (2020) for further details.

Mice are trained to detect of a static visual grating of varying contrast (a Gabor patch) in either the left or right visual field (Figure 1A). The visual stimulus is coupled with the movements of a response wheel, and animals indicate their choices by turning the wheel left or right to bring the grating to the center of the screen (Burgess et al., 2017). The visual stimulus appears on the screen after an auditory “go cue” indicates the start of the trial and only if the animal holds the wheel for 0.2-0.5 sec. Correct decisions are rewarded with sweetened water (10% sucrose solution, Guo et al. (2014)), while incorrect decisions are indicated by a noise burst and are followed by a longer inter-trial interval (2 seconds).

Mice begin training on a “basic” version of the task, where the probability of a stimulus appearing on the left or the right is 50:50. Training begins with a set of “easy” contrasts (100% and 50%), and harder contrasts (25%, 12.5%, 6.25%, and 0%) are introduced progressively according to predefined performance criteria. After a mouse achieves a predefined performance criteria, a “biased” version of the task is introduced where probability switches in blocks of trials between 20:80 favoring the right and 80:20 favoring the left.

#### Akrami Task

Here we review the relevant features of the task, as well as the rat and human subject training protocols from the Akrami task. Please refer to Akrami et al. (2018) for further details.

Rats were trained on an auditory delayed comparison task, adapted from a tactile version (Fassihi et al., 2014). Training occurred within three-port operant conditioning chambers, in which ports are arranged side-by-side along one wall, with two speakers placed above the left and right nose ports. Figure 5A shows the task structure. Rat subjects initiate a trial by inserting their nose into the centre port, and must keep their nose there (fixation period), until an auditory “go” cue plays. The subject can then withdraw and orient to one of the side ports in order to receive a reward of water. During the fixation period, two auditory stimuli, Tones A and B, separated by a variable delay, are played for 400 ms, with short delay periods of 250 ms inserted before Tone A and after Tone B. The stimuli consist of broadband noise (2,000–20,000 Hz), generated as a series of sound pressure level (SPL) values sampled from a zero-mean normal distribution. The overall mean intensity of sounds varies from 60–92 dB. Rats should judge which out of the two stimuli, Tones A and B, had the greater SPL standard deviation. If Tone A *>* B, the correct action is to poke the nose into the right-hand nose port in order to collect the reward, and if Tone A *<* B, rats should orient to the left-hand nose port.

Trial durations are independently varied on a trial-by-trial basis, by varying the delay interval between the two stimuli, which can be as short as 2s or as long as 12s. Rats progressed through a series of shaping stages before the final version of the delayed comparison task, in which they learned to: associate light in the centre poke with the availability of trials; associate special sounds from the side pokes with reward; maintain their nose in the centre poke until they hear an auditory “go” signal; and compare the Tone A and B stimuli. Training began when rats were two months old, and typically required three to four months for rats to display stable performance on the complete version of the task.

In the human version of the task, similar auditory stimuli to those used for rats were used (see Figure 7A). Subjects received, in each trial, a pair of sounds played from ear-surrounding noise-cancelling headphones. The subject self-initiated each trial by pressing the space bar on the keyboard. Tone A was then presented together with a green square on the left side of a computer monitor in front of the subject. This was followed by a delay period, indicated by “WAIT!” on the screen, then Tone B was presented together with a red square on the right side of the screen. At the end of the second stimulus and after the go cue, subjects were required to compare the two sounds and decide which one was louder, then indicate their choice by pressing the “k” key with their right hand (Tone B was louder) or the “s” key with their left hand (Tone A was louder). Written feedback about the correctness of their response was provided on the screen, for individual trials as well as the average performance updated every ten trials.

### QUANTIFICATION AND STATISTICAL ANALYSIS

#### Cross-validation Procedure

When making predictions about specific trials, the model should not be trained using those trials. We implement a 10-fold cross-validation procedure where the model is fit using a training set composed of a random 90% of trials at a time, and the remaining 10% is used for predicting and testing. We modify the prior Σ such that the gaps created by removing the 10% of test set trials are taken into account. For example, if trial *t* is in the test set and trials *t*− 1 and *t*+1 are in the training set, then we modify the value on the diagonal of Σ corresponding to trial *t* −1 from *σ*^2^ to 2*σ*^2^ to account for the missing entry in Σ created by omitting trial *t* from the training set.

To predict the animal’s choice at a test trial *t*, we first infer the weights for a training set of trials that excludes *t*, as described above. Then we approximate **w**_*t*_ by interpolating from the nearest adjacent trials in the training set. We repeat this to obtain a set of *predicted weights* for each trial. Using the predicted weights **w**_*t*_ and the input vector **x**_*t*_ for that trial, we can calculate a predicted choice probability *P* (Go Right) (as in Figure 5), and compare this to the actual choice *y*_*t*_ to calculate the predicted accuracy (as in Figure 8; see below for more details). Using the predicted weights to calculate a cross-validated log-likelihood *L* can be used to choose between models in lieu of approximate log-evidence.

#### Calculation of Posterior Credible Intervals

In order to estimate the extent to which our recovered weights **w** are constrained by the data, we calculate a posterior credible interval over the time-varying weight trajectories (e.g., the shaded regions shown in Figure 5B). Specifically, we approximate the 95% posterior credible interval by using 1.96 standard deviations. The standard deviation is calculated by taking the square-root of the diagonal of the covariance matrix at **w**_MAP_.

The covariance matrix can be approximated by the inverse Hessian, but inversion of a large matrix can be challenging. Here we adapt a fast method for inverting block-tridiagonal matrices (Rybicki and Hummer, 1991), taking advantage of the fact that our Hessian (while extremely large) has a block-tridiagonal structure, and that we only need the diagonal of the inverse Hessian. If *H* is the Hessian matrix of our weights at the posterior peak **w**_MAP_ (also with the highest log-evidence, according to the optimization procedure in Algorithm 1), this method calculates a diagonal matrix,

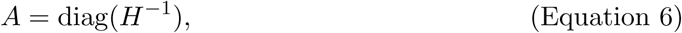

such that we can take 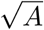 to estimate one standard deviation for each weight on each trial. The algorithm requires order *TK*^3^ scalar operations for calculating the central blocks of our inverse Hessian (for *T* trials and *K* weights).

To calculate the posterior credible intervals for our hyperparameters *θ*, we take the same approach of inverting the *K* × *K* hyperparameter Hessian matrix (or 2*K* × 2*K* if *σ*_day_ is used). The difficulty here is not in inverting this Hessian matrix, since it is much smaller, but in calculating the Hessian in the first place. We calculate each entry in our Hessian manually, numerically determining each of the *K*(*K* + 1)*/*2 unique entries. Once we have determined the Hessian matrix for the hyperparameters, it is straightforward to calculate the 95% posterior credible intervals for each hyperparameter, using the same procedure as in the case for the weights.

#### Calculation of Predicted and Empirical Measures

Here we explain how the accuracy and bias measures (Figure 5C and D) were calculated. Accuracy is *empirically* calculated by constructing a vector of length *T* where the *i*^th^ entry is a 0 if the animal answered incorrectly on the *i*^th^ trial and a 1 if answered correctly. This vector is then smoothed (we use a Gaussian kernel of *σ* = 50) to get the *empirical* accuracy plotted in pink in Figure 5C, with a 95% credible interval in shaded pink. To calculate the *predicted* accuracy, we use the cross-validation procedure detailed above to calculate *P* (Correct Answer) for each trial which we then smooth with the same Gaussian kernel to get the black line in Figure 5C.

As a simple way of characterizing only the *task-irrelevant* tendency to make a specific choice as the bias, here we define empirical bias as a preference for left or right only on *incorrect* trials. Specifically, we construct a vector of length *T* where the *i*^th^ entry is the animal’s choice minus the correct answer in the *i*^th^ trial, where both choice and answer are coded such that “left” is 0 and “right” is 1. Thus, the *empirical* bias on each trial is one of {−1, 0, +1}. This vector is then smoothed (again using a Gaussian kernel of *σ* = 50) to get the empirical bias plotted in purple in Figure 5D, with a 95% credible interval in shaded purple. We calculate the *predicted* bias in a similar manner to the predicted accuracy, using cross-validated weights to calculate *P* (Right) for each trial and substituting that value in for the animal’s choice — thus for each trial we get a value from a continuous interval (−1, +1) which we then smooth with the same Gaussian kernel to get the black line in Figure 5C.

## DATA AND CODE AVAILABILITY

Our code for fitting psychophysical weights to behavioral data is distributed as a GitHub repository (under a MIT license): https://github.com/nicholas-roy/psytrack. This code is also made easily accessible as a Python package, PsyTrack (installed via `pip install psytrack`). Our Python package relies on the standard SciPy scientific computing libraries as well as the Open Neurophysiology Environment produced by the IBL (Jones et al., 2001; Hunter, 2007; Bonacchi et al., 2019). All the data analyzed from IBL et al. (2020) and Akrami et al. (2018) is also publicly available (see the Key Resources Table for links).

We have assembled a Google Colab notebook that will automatically download the raw data and allow precise reproduction of all figures from the paper. Our analyses can be easily extended to additional experimental subjects and hopefully act as a template for application of PsyTrack to new datasets.

---

**Figure S1.**
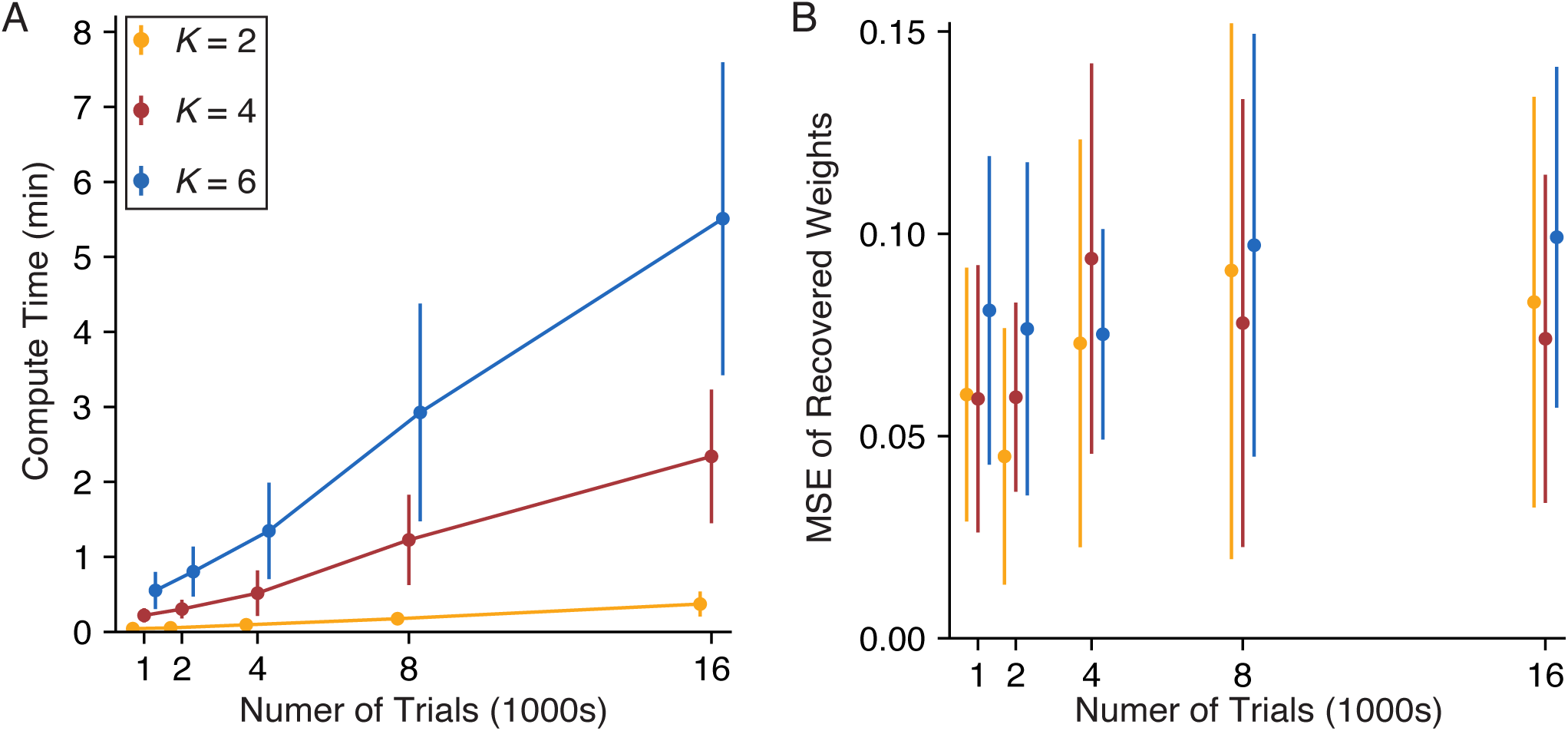
Related to Figure 2, Compute time and model accuracy. **(A)** The model fitting time as a function of the number of weights *K* = {2, 4, 6} and the number of trials *T* = {1000, 200, 4000, 8000, 16000}. Weights are simulated as in Figure 2A. For each pair of *K* weights and *T* trials, 20 sets of weights are randomly simulated (with each log_2_(*σ*_*k*_) ∼ 𝕌(−7.5, −3.5); no *σ*_day_ was used), and recovered by the model (the calculation of credible intervals on the weights was omitted). We plot the average recovery time across the 20 iterations, ±1 standard deviation. We can see that even a reasonably complex model (6 weights and 16000 trials) only takes around 5 minutes to fit on average. All models were fit on a 2012 MacBook Pro with a 2.3 GHz Quad-Core Intel i7 processor. **(B)** The mean-squared error (MSE) calculated across all weights across all trials, as a function of the total number of weights and total number of trials in the model. Calculations used the same models run in (A). The average MSE across the 20 iterations of each model is plotted, ±1 standard deviation. We can see that the recovery of the weights is relatively independent of the number of weights (color coded as in (A)) and number of trials.

**Figure S2.**
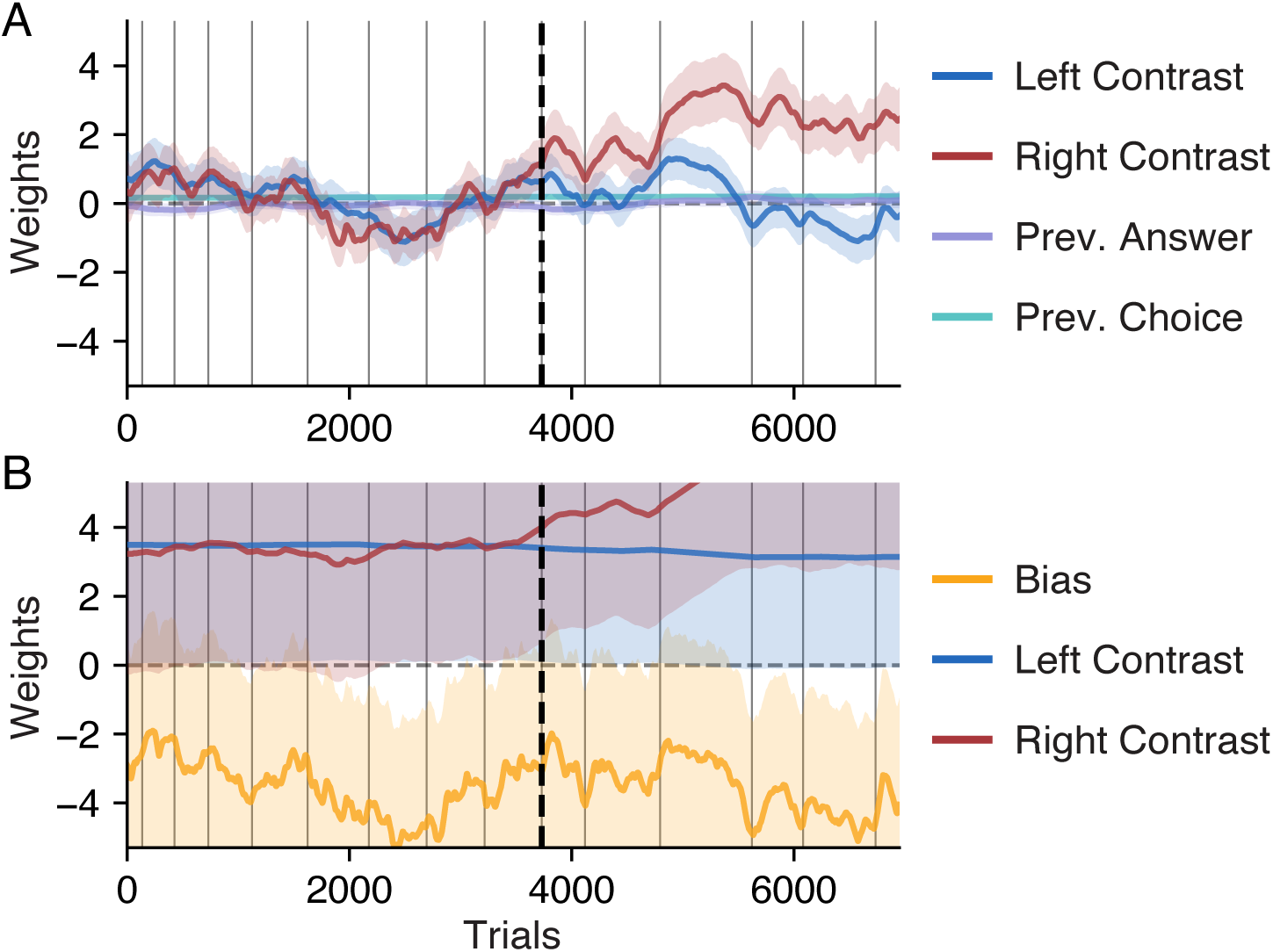
Related to Figure 3, Adding weights to early training sessions in IBL mice. **(A)** Here we refit the data first presented in Figure 3B, but now adding two additional weights: a Previous (Correct) Answer and Previous Choice weight. While these history regressor weights massively impact choice behavior during early training in all the Akrami Rats (see Figure 6D and E), they have negligible impact on the behavior of our example IBL mouse. **(B)** Here we refit the data first presented in Figure 3B, but now adding a bias weight. While a bias weight is very useful for describing the behavior of IBL mice later in training (see Figure 4), we omit the bias weight during early training due to an issue of non-identifiability. During the initial sessions of training, IBL mice are only presented with “easy” contrast values of 50% and 100%. These contrasts are perceptually very similar (i.e. a 100% contrast is not twice as difficult as a 50% contrast), which we account for with a tanh transformation of the contrasts (see Figure S3). Thus, the task in the earliest stages of training has effectively only two types of trials: ∼ 100% left contrast trials and ∼ 100% right contrast trials. With a task this simple, behavior is over-parameterized by having a bias weight and two contrast weights, introducing the non-identifiability. Explicitly, a model with hypothetical weight values of [bias, left, right] = [0, -1, +1] is nearly identical to a model with values [-1, 0, 2]; in fact, there are an infinite number of weight values for these three weights that would all describe behavior on this simplified task in approximately the same way. Fortunately, the posterior credible intervals on the weights can indicate that a model is in a non-identifiable regime. Because there are so many settings of possible weight value, the intervals become abnormally large and overlapping, as shown here. See the Methods for more details about model non-identifiability.

**Figure S3.**
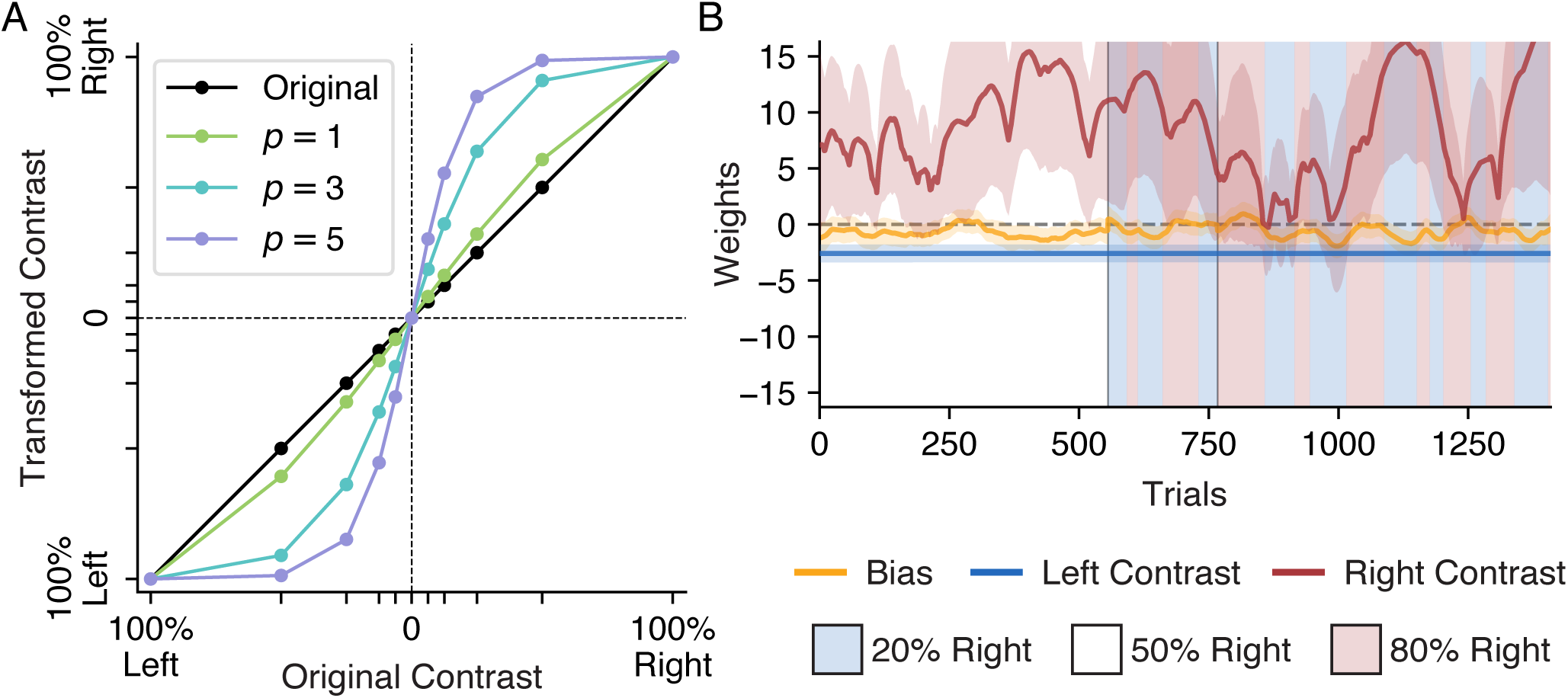
Related to Figure 4, The impact of the tanh transformation of IBL contrasts on model weights. **(A)** The effect of the tanh transformation on the IBL contrast values for several settings of the free parameter *p*. A tanh transformation is applied to the contrast values, *c*, in the IBL task such that the relative values of the transformed contrasts, *ĉ*_*p*_ = tanh(*pc*)*/* tanh(*p*), better aligns with their relative perceptual difficulty. In this work, we use *p* = 5 to transform the contrasts (purple line), such that *c* = {0, 0.0625, 0.125, 0.25, 0.5, 1} are transformed to *ĉ*_5_ = {0, 0.303, 0.555, 0.848, 0.987, 1} (left contrasts are coded as having negative value here). This value was anecdotally observed to work well across a large variety of mice and sessions, but it could be optimized for each model. **(B)** Here we refit the data first presented in Figure 4B, forgoing the tanh transformation and using the original contrast values *c* instead. We see that the right contrast weight grows to massive values and fluctuates wildly. Using Equation 1, we can calculate that with a weight of 15, the model predicts that a 100% right contrast would result in a *P* (Go Right) of over 99.9999% (disregarding the impact of the much smaller bias weight, for simplicity). This is an absurdly confident prediction, even for the best trained mouse, but it represents a compromise the model was forced to make. We can calculate that the predicted *P* (Go Right) on the most difficult right contrast value, a 6.25% contrast, is a much more reasonable 71.9%. A well-trained mouse could certainly be performing better than this on the hardest contrast. However, in order to reflect a higher *P* (Go Right) on this hard contrast, the right contrast weight would need to become even greater, resulting in even more absurdly confident predictions on the 100% contrast trials. All right contrast values share the same weight, forcing a single compromise weight value between them. By using the raw contrast value, the model assumes that a 100% right contrast is 16 times more salient to the mouse than a 6.25% contrast, which is empirically untrue (Busse et al., 2011). Applying the tanh transformation brings the relative value of the two contrasts into a much more reasonable regime. With *p* = 5, a 6.25% contrast is encoded as 0.303 while the 100% contrast remains at 1.0, making the 100% contrast only 3 time more salient than the 6.25% contrast. See the Methods for more details about the parametrization of input variables.

**Figure S4.**
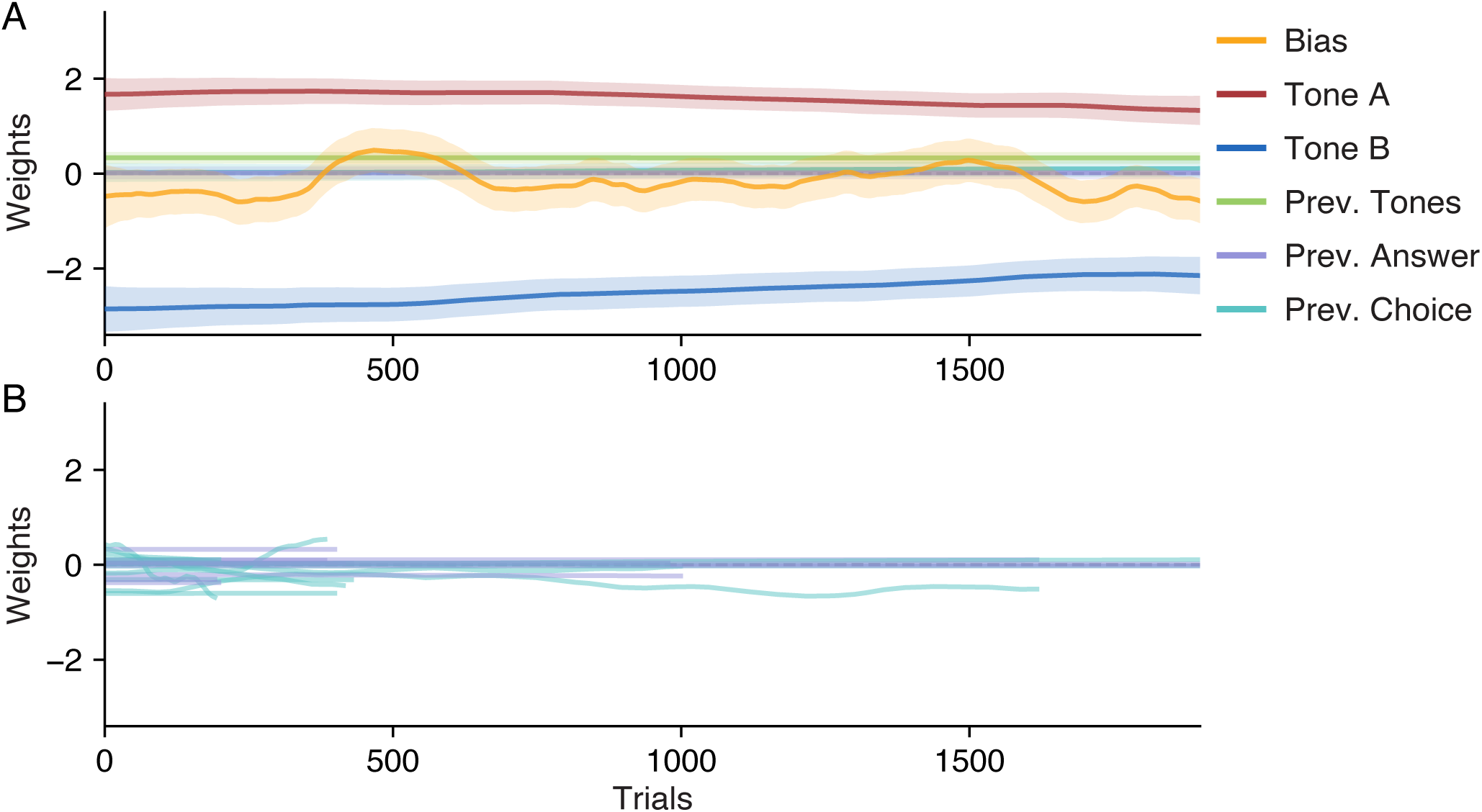
Related to Figure 7, Modeling the Akrami human subjects with the Previous Choice and Previous Answer weights. **(A)** Here we refit the data presented in Figure 7B, but now we add both a Previous Choice and a Previous (Correct) Answer weight. Since human subjects are given task instructions before starting, we would not expect their behavior to be affected by either their choice or the correct answer on the previous trial. Indeed, we see that both new history regressor weights are always approximately 0 for our example subject. **(B)** Here we refit the data presented in Figure 7C, but now we add both a Previous Choice and a Previous (Correct) Answer weight for all the human subjects. We plot only the two new weights from each refit model, for clarity. We see that, in general, both the choice and the correct answer on the previous trial have a negligible impact on human choice behavior.

## References

Akrami, A., Kopec, C.D., Diamond, M.E., Brody, C.D., 2018. Posterior parietal cortex represents sensory history and mediates its effects on behaviour. Nature 554, 368.

Bak, J.H., Choi, J.Y., Akrami, A., Witten, I., Pillow, J.W., 2016. Adaptive optimal training of animal behavior, in: Advances in Neural Information Processing Systems, pp. 1947–1955.

Bak, J.H., Pillow, J.W., 2018. Adaptive stimulus selection for multi-alternative psychometric functions with lapses. Journal of Vision 18, 4. URL: +http://dx.doi.org/10.1167/18.12.4, doi: 10.1167/18.12.4, /data/journals/jov/937613/i1534-7362-18-12-4.pdf.

Bishop, C.M., 2006. Pattern recognition and machine learning. Springer.

Bonacchi, N., Chapuis, G., Churchland, A., Harris, K.D., Rossant, C., Sasaki, M., Shen, S., Steinmetz, N.A., Walker, E.Y., Winter, O., et al., 2019. Data architecture and visualization for a large-scale neuroscience collaboration. BioRxiv, 827873.

Brunton, B.W., Botvinick, M.M., Brody, C.D., 2013. Rats and humans can optimally accumulate evidence for decision-making. Science 340, 95–98. URL: https://doi.org/10.1126/science.1233912, doi: 10.1126/science.1233912.

Burgess, C.P., Lak, A., Steinmetz, N.A., Zatka-Haas, P., Reddy, C.B., Jacobs, E.A., Linden, J.F., Paton, J.J., Ranson, A., Schröder, S., et al., 2017. High-yield methods for accurate two-alternative visual psychophysics in head-fixed mice. Cell reports 20, 2513–2524.

Busse, L., Ayaz, A., Dhruv, N.T., Katzner, S., Saleem, A.B., Schölvinck, M.L., Zaharia, A.D., Carandini, M., 2011. The detection of visual contrast in the behaving mouse. Journal of Neuroscience 31, 11351–11361. URL: http://jneurosci.org/content/31/31/11351, doi: 10.1523/JNEUROSCI.6689-10.2011, arXiv:http://jneurosci.org/content/31/31/11351.full.pdf.

Carandini, M., 2012. From circuits to behavior: a bridge too far? Nature Neuroscience 15, 507–509. URL: https://doi.org/10.1038/nn.3043, doi: 10.1038/nn.3043.

Carandini, M., Churchland, A.K., 2013. Probing perceptual decisions in rodents. Nature neuroscience 16, 824.

Churchland, A.K., Kiani, R., Shadlen, M.N., 2008. Decision-making with multiple alternatives. Nature Neuroscience 11, 693–702. URL: https://doi.org/10.1038/nn.2123, doi: 10.1038/nn.2123.

Cohen, Y., Schneidman, E., 2013. High-order feature-based mixture models of classification learning predict individual learning curves and enable personalized teaching. Proceedings of the National Academy of Sciences 110, 684–689.

Corrado, G.S., Sugrue, L.P., Seung, H.S., Newsome, W.T., 2005. Linear-nonlinear-poisson models of primate choice dynamics. J Exp Anal Behav 84, 581–617.

Courville, A.C., Daw, N.D., Touretzky, D.S., 2006. Bayesian theories of conditioning in a changing world. Trends in cognitive sciences 10, 294–300.

Daw, N.D., Doya, K., 2006. The computational neurobiology of learning and reward. Current opinion in neurobiology 16, 199–204.

Dayan, P., Balleine, B.W., 2002. Reward, motivation, and reinforcement learning. Neuron 36, 285–298.

Duncker, L., Sahani, M., 2018. Temporal alignment and latent gaussian process factor inference in population spike trains, in: Advances in Neural Information Processing Systems, pp. 10445–10455.

Fassihi, A., Akrami, A., Esmaeili, V., Diamond, M.E., 2014. Tactile perception and working memory in rats and humans. Proceedings of the National Academy of Sciences 111, 2331–2336.

Frund, I., Wichmann, F.A., Macke, J.H., 2014. Quantifying the effect of intertrial dependence on perceptual decisions. Journal of Vision 14, 9–9. URL: https://doi.org/10.1167/14.7.9, doi: 10.1167/14.7.9.

Gao, Y., Archer, E.W., Paninski, L., Cunningham, J.P., 2016. Linear dynamical neural population models through nonlinear embeddings, in: Advances in Neural Information Processing Systems, pp. 163–171.

Green, D.M., Swets, J.A., 1966. Signal Detection Theory and Psychophysics. Wiley, New York.

Guo, Z.V., Hires, S.A., Li, N., O’Connor, D.H., Komiyama, T., Ophir, E., Huber, D., Bonardi, C., Morandell, K., Gutnisky, D., et al., 2014. Procedures for behavioral experiments in head-fixed mice. PloS one 9.

Hunter, J.D., 2007. Matplotlib: A 2d graphics environment. Computing in science & engineering 9, 90–95.

IBL, Aguillon-Rodriguez, V., Angelaki, D.E., Bayer, H.M., Bonacchi, N., Carandini, M., Cazettes, F., Chapuis, G.A., Churchland, A.K., Dan, Y., Dewitt, E.E., Faulkner, M., Forrest, H., Haetzel, L.M., Hausser, M., Hofer, S.B., Hu, F., Khanal, A., Krasniak, C.S., Laranjeira, I., Mainen, Z.F., Meijer, G.T., Miska, N.J., Mrsic-Flogel, T.D., Murakami, M., Noel, J.P., Pan-Vazquez, A., Sanders, J.I., Socha, K.Z., Terry, R., Urai, A.E., Vergara, H.M., Wells, M.J., Wilson, C.J., Witten, I.B., Wool, L.E., Zador, A., 2020. A standardized and reproducible method to measure decision-making in mice. bioRxiv URL: https://www.biorxiv.org/content/early/2020/01/19/2020.01.17.909838, doi: 10.1101/2020.01.17.909838, arXiv:https://www.biorxiv.org/content/early/2020/01/19/2020.01.17.909838.full.pdf.

Jones, E., Oliphant, T., Peterson, P., et al., 2001. SciPy: Open source scientific tools for Python. URL: http://www.scipy.org/.

Kattner, F., Cochrane, A., Green, C.S., 2017. Trial-dependent psychometric functions accounting for perceptual learning in 2-afc discrimination tasks. Journal of vision 17, 3–3.

Kobak, D., Brendel, W., Constantinidis, C., Feierstein, C.E., Kepecs, A., Mainen, Z.F., Qi, X.L., Romo, R., Uchida, N., Machens, C.K., 2016. Demixed principal component analysis of neural population data. Elife 5, e10989.

Krakauer, J.W., Ghazanfar, A.A., Gomez-Marin, A., MacIver, M.A., Poeppel, D., 2017. Neuroscience needs behavior: correcting a reductionist bias. Neuron 93, 480–490.

Licata, A.M., Kaufman, M.T., Raposo, D., Ryan, M.B., Sheppard, J.P., Churchland, A.K., 2017. Posterior parietal cortex guides visual decisions in rats. Journal of Neuroscience 37, 4954–4966.

Lu, Z., Williamson, S., Kaufman, L., 1992. Behavioral lifetime of human auditory sensory memory predicted by physiological measures. Science 258, 1668–1670.

Meshulam, L., Gauthier, J.L., Brody, C.D., Tank, D.W., Bialek, W., 2019. Coarse graining, fixed points, and scaling in a large population of neurons. Physical review letters 123, 178103.

Murphy, R.A., Mondragon, E., Murphy, V.A., 2008. Rule learning by rats. Science 319, 1849–1851. URL: https://doi.org/10.1126/science.1151564, doi: 10.1126/science.1151564.

Nocedal, J., Wright, S.J., 2006. Quasi-newton methods. Numerical optimization, 135–163.

O’Doherty, J.P., Dayan, P., Friston, K., Critchley, H., Dolan, R.J., 2003. Temporal difference models and reward-related learning in the human brain. Neuron 38, 329–337.

Papadimitriou, C., Ferdoash, A., Snyder, L.H., 2015. Ghosts in the machine: memory interference from the previous trial. Journal of neurophysiology 113, 567–577.

Piet, A.T., Hady, A.E., Brody, C.D., 2018. Rats adopt the optimal timescale for evidence integration in a dynamic environment. Nature Communications 9. URL: https://doi.org/10.1038/s41467-018-06561-y, doi: 10.1038/s41467-018-06561-y.

Pisupati, S., Chartarifsky-Lynn, L., Khanal, A., Churchland, A.K., 2019. Lapses in perceptual decisions reflect exploration. bioRxiv, 613828.

Prerau, M.J., Smith, A.C., Eden, U.T., Kubota, Y., Yanike, M., Suzuki, W., Graybiel, A.M., Brown, E.N., 2009. Characterizing learning by simultaneous analysis of continuous and binary measures of performance. Journal of neurophysiology 102, 3060–3072.

Roy, N.A., Bak, J.H., Akrami, A., Brody, C., Pillow, J.W., 2018a. Efficient inference for time-varying behavior during learning, in: Advances in neural information processing systems, pp. 5695–5705.

Roy, N.A., Bak, J.H., Pillow, J.W., 2018b. PsyTrack: Open source dynamic behavioral fitting tool for Python. URL: https://github.com/nicholas-roy/psytrack.

Rybicki, G.B., Hummer, D.G., 1991. An accelerated lambda iteration method for multilevel radiative transfer. I-Non-overlapping lines with background continuum; Appendix B. Astronomy and Astrophysics 245, 171–181.

Sahani, M., Linden, J.F., 2003. Evidence optimization techniques for estimating stimulus-response functions, in: Advances in neural information processing systems, pp. 317–324.

Schultz, W., Dayan, P., Montague, P.R., 1997. A neural substrate of prediction and reward. Science 275, 1593–1599.

Semedo, J.D., Zandvakili, A., Machens, C.K., Byron, M.Y., Kohn, A., 2019. Cortical areas interact through a communication subspace. Neuron 102, 249–259.

Smith, A.C., Frank, L.M., Wirth, S., Yanike, M., Hu, D., Kubota, Y., Graybiel, A.M., Suzuki, W.A., Brown, E.N., 2004. Dynamic analysis of learning in behavioral experiments. Journal of Neuroscience 24, 447–461.

Sutton, R.S., 1988. Learning to predict by the methods of temporal differences. Machine learning 3, 9–44.

Sutton, R.S., Barto, A.G., 2018. Reinforcement learning: An introduction. MIT press.

Suzuki, W.A., Brown, E.N., 2005. Behavioral and neurophysiological analyses of dynamic learning processes. Behavioral and cognitive neuroscience reviews 4, 67–95.

Tajima, S., Drugowitsch, J., Patel, N., Pouget, A., 2019. Optimal policy for multi-alternative decisions. Nature neuroscience 22, 1503–1511.

Usher, M., Tsetsos, K., Yu, E.C., Lagnado, D.A., 2013. Dynamics of decision-making: from evidence accumulation to preference and belief. Frontiers in Psychology 4. URL: https://doi.org/10.3389/fpsyg.2013.00758, doi: 10.3389/fpsyg.2013.00758.

Williams, A.H., Poole, B., Maheswaranathan, N., Dhawale, A.K., Fisher, T., Wilson, C.D., Brann, D.H., Trautmann, E.M., Ryu, S., Shusterman, R., et al., 2020. Discovering precise temporal patterns in large-scale neural recordings through robust and interpretable time warping. Neuron 105, 246–259.

Wu, A., Roy, N.A., Keeley, S., Pillow, J.W., 2017. Gaussian process based nonlinear latent structure discovery in multivariate spike train data, in: Advances in Neural Information Processing Systems, pp. 3499–3508.

